# A global atlas of deep-sea cold seep viruses uncovers extensive genomic novelty and biotechnological potential

**DOI:** 10.64898/2025.12.10.692286

**Authors:** Xinyue Liu, Xiaomei Guo, Lan Xue, Zijian Lu, Jieni Wang, Jing Liao, Lu Liu, Yanbin Chen, Yongyi Peng, Yingchun Han, Fabai Wu, Rui Cheng, Xiyang Dong

**Author notes:** Correspondence:* *(X.D.);* *(R.C.). These authors contributed equally to this work (Xinyue Liu, Xiaomei Guo, Lan Xue).

## Abstract

Cold seeps are globally distributed deep-sea ecosystems harboring diverse chemosynthetic microbial communities, yet their viral components remain poorly understood. Here, we establish the Global Cold Seep Virome, a comprehensive catalog of 191,937 species-level viral operational taxonomic units (vOTUs) and 2.44 million viral protein clusters reconstructed from 314 metagenomes spanning 18 cold seep sites worldwide. Most vOTUs are distinct from those in existing environmental viromes, suggesting that cold seeps harbor a largely unexplored viral reservoir. Host prediction suggests diverse viral associations with key prokaryotes, including anaerobic methanotrophic archaea. Protein annotation reveals that these viruses encode widespread uncharacterized proteins and viral hallmark proteins with divergent physicochemical properties, potentially contributing to their ecological adaptability. We also identify putative auxiliary metabolic genes, anti-defense genes, and lysins, and experimentally validate representative anti-defense proteins and lysins with strong activities. This resource expands current knowledge of deep-sea viral diversity and provides a foundation for studying virus-host interactions, testing biogeochemical hypotheses, and exploring biotechnological potential in cold seep ecosystems.

## Introduction

Viruses are central components of microbial ecosystems, shaping host population dynamics, mediating gene exchange, and influencing metabolic fluxes across diverse environments^1,2^. Despite major advances in marine virology and other accessible environments^3–6^, much of the global virosphere remains poorly characterized, particularly in the deep sea where limited sampling and extreme physical constraints restrict our understanding^7^. Metagenomic surveys of deep-sea ecosystems, including hydrothermal vents, seamounts, and hadal trenches, consistently reveal high levels of viral novelty, with most genes lacking detectable homology to known sequences^8–12^. These observations suggest that deep-sea environments harbor distinct viral lineages and functions, yet the ecological roles and molecular strategies of these communities remain largely unresolved due to incomplete genome recovery and fragmented functional annotation.

Cold seeps provide a tractable setting for investigating deep-sea microbial and viral processes. The sustained release of methane and other hydrocarbons supports dense chemosynthetic communities dominated by archaeal and bacterial groups that mediate methane oxidation, sulfur and nitrogen transformations, and organic carbon turnover^13,14^. These processes are structured by steep geochemical gradients and relatively stable physical conditions, features that distinguish cold seeps from the more dynamic environments typically profiled in viral ecology. Although previous studies indicate that cold seep sediments contain diverse and taxonomically novel viruses^15,16^, their interactions with host physiology, responses to sediment geochemistry, and potential contributions to major metabolic pathways remain insufficiently characterized. Likewise, the functional potential of cold seep viruses, including auxiliary metabolic genes (AMGs)^17^, immune antagonists such as anti-defense proteins and lysins^18–20^, and enzymes adapted to low temperature and high hydrostatic pressure, has not been systematically explored. Mining this virosphere may yield cold-adapted enzymes with value in biotechnology and antimicrobial development^21–23^.

Progress toward addressing these questions has been limited by two major constraints: the difficulty of reconstructing complete viral genomes from complex sediments and linking them to microbial hosts, and the high proportion of viral genes (40-90%) that remain functionally unannotated^24^. Recent advances in long-read metagenomics and genome-resolved host prediction now enable more complete viral genome recovery and more confident host associations^25,26^. In parallel, structure-guided and large language model-based annotation methods have improved functional inference for divergent viral proteins that typically evade conventional sequence-based approaches^27^. These developments provide an opportunity to more fully resolve the genetic and biochemical landscape of deep-sea viromes.

In this study, we assemble the Global Cold Seep Virome (GCSV), a comprehensive viral genome and protein resource derived from metagenomic datasets spanning multiple cold seep systems worldwide. By integrating viral genome assemblies, host-associated microbial genomes, and an extensive viral protein catalog, we characterize the taxonomic diversity, functional potential, and evolutionary features of cold seep viruses at a broad geographic scale. This framework provides a basis for generating testable hypotheses about virus-host interactions, potential viral contributions to biogeochemical processes, and the molecular features of deep-sea-derived viral proteins. We further validate representative viral enzymes to demonstrate the utility of this resource for functional and biotechnological exploration. Together, the GCSV establishes a curated reference for deep-sea viral ecology and provides a foundation for future mechanistic and applied investigations.

## Results

### The largest cold seep virome with high novelty and genomic stability

By mining metagenomic data from 314 samples collected across 18 globally distributed cold seeps (**Fig. 1a and Table S1; Methods and materials**), we constructed the GCSV, which comprises 38 bottom-water short-read metagenomes, 269 sediment short-read metagenomes, and seven sediment long-read metagenomes. In total, the dataset contains 0.5 Tb of PacBio HiFi long-read data and 10 Tb of Illumina short-read data, representing a > 25-fold increase in data volume compared to the 0.38 Tb available in our previous study^15^. These data support the GCSV as the most comprehensive cold seep viral catalog generated to date^15,16,28^ and one of the largest deep-sea viral resources available^7^.

**Figure 1.**
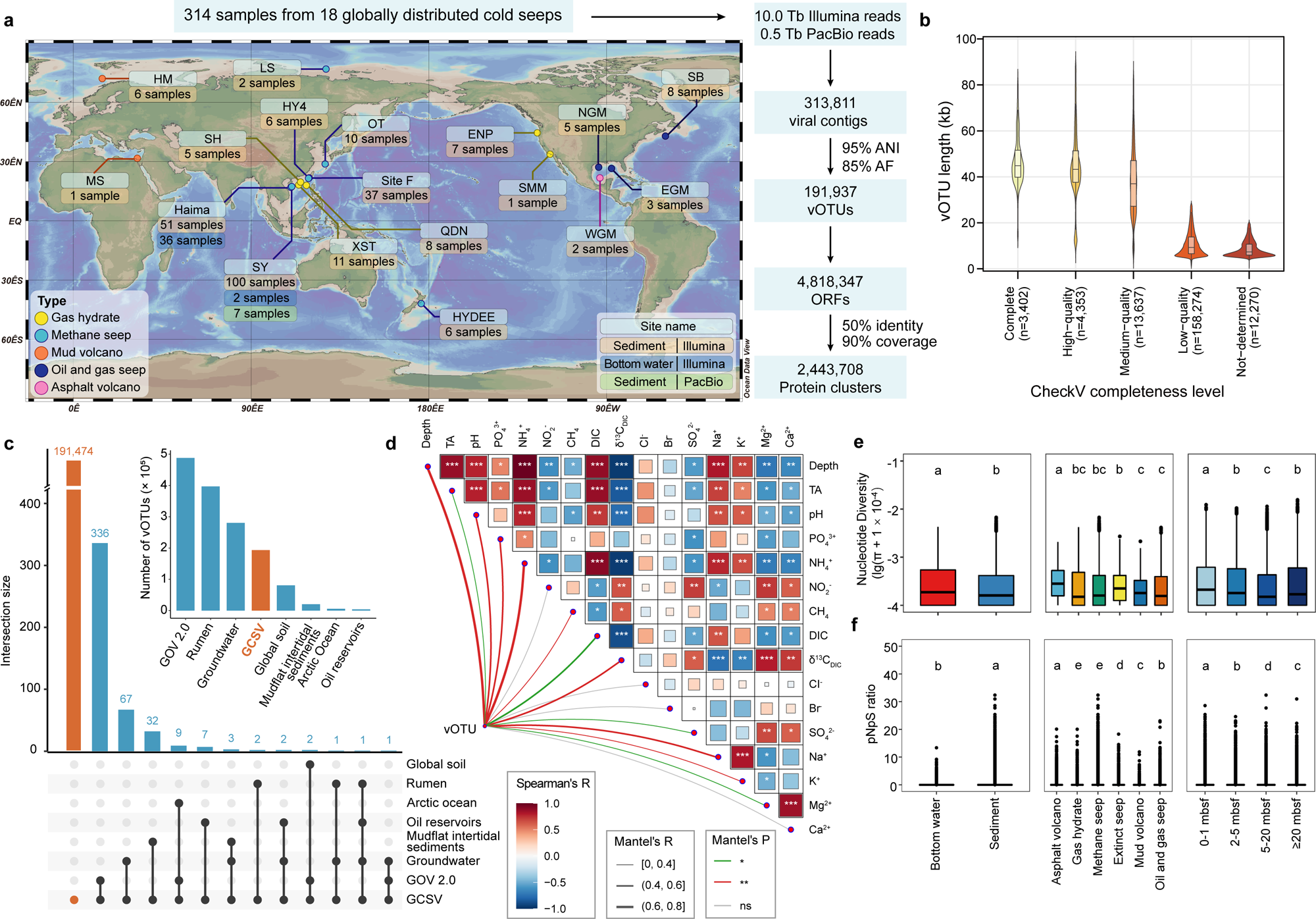
Overview and diversity of the Global Cold Seep Virome (GCSV). **(a)** Geographic distribution of 314 samples collected from 18 globally distributed cold seeps and summary of dataset size across major processing steps (see **Methods and materials, Fig. S1** and **Table S1**). ANI, average nucleotide identity; AF, alignment fraction; vOTU, viral operational taxonomic unit; ORF, open reading frame. **(b)** Distribution of viral genome lengths across CheckV quality tiers. Violin plots and boxplots show length variation of vOTUs categorized by CheckV estimates (**Table S2**). **(c)** Shared and unique vOTUs between the GCSV and other viral databases. The UpSet plot (bottom left) displays the number of vOTUs shared between GCSV and other datasets, with GCSV-unique vOTUs highlighted in orange. The bar plot (upper right) shows the total number of vOTUs in each database. **(d)** Correlation between vOTU abundance and environmental parameters. In the lower-left panel, links represent results of Mantel’s tests. Line width corresponds to correlation strength (*R* value), and colors indicate statistical significance (*P* value). In the upper-right panel, Spearman correlation coefficients among environmental parameters are shown. Colors represent Spearman’s *R*, and asterisks denote significance levels (*, *P* < 0.05; **, *P* < 0.01; ***, *P* < 0.001). Analyses were based on 20 samples (P1-P20) from site W07. Detailed statistical summaries are provided in **Table S17**. **(e)** Nucleotide diversity (π) of cold seep viruses across sample types, seep types, and sediment depths. **(f)** Gene-level selection patterns assessed by pN/pS ratios across sample types, seep types, and sediment depths. Letters above boxplots denote statistical groupings based on pairwise Wilcoxon rank-sum tests with Benjamini-Hochberg correction (*P* < 0.05). Groups labeled with different letters show statistically significant differences. mbsf, meters below seafloor. Detailed statistical summaries are provided in **Table S3**.

Using three complementary virus identification pipelines followed by systematic quality control (**Figs. S1-S2; Methods and materials**), we identified 313,811 viral contigs and clustered them at 95% average nucleotide identity (ANI) and 85% alignment fraction (AF). This procedure yielded 191,937 non-redundant viral operational taxonomic units (vOTUs), approximating species-level viral genomes, including 12,226 proviruses (**Fig. 1a and Table S2**). CheckV assessment revealed 3,402 complete, 4,353 high-quality, and 13,637 medium-quality genomes, together accounting for 11.15% of the GCSV catalog (**Fig. S3a**). Average genome length scaled with quality tier: high-quality (56.63 kb) and complete (53.40 kb) genomes were substantially longer than medium-quality (44.04 kb), low-quality (12.94 kb), and undetermined-quality vOTUs (10.09 kb) (**Fig. 1b**). A total of 71,530 vOTUs (37.27% of the entire catalog) originated from PacBio assemblies, which exhibited larger mean genome size (19.35 kb) and a higher proportion of genomes with ≥ 50% estimated completeness (12.14%) compared with Illumina-derived vOTUs (15.10 kb; 10.55%). These results underscore the advantages of long-read sequencing for reconstructing longer and more complete viral genomes. To evaluate habitat specificity and global novelty, we compared the GCSV against viral datasets from diverse ecosystems, including mudflat intertidal sediments^29^, soils^30^, groundwater^6^, the Global Ocean Virome 2 (GOV 2.0)^4^, the Arctic Ocean surface microlayer^31^, deep subsurface oil reservoirs^5^, and the rumen microbiome^32^ **(Fig. 1c)**. Remarkably, only 0.24% of GCSV vOTUs showed species-level overlap with viruses from these environments, highlighting the high novelty and strong habitat specificity of cold seep viral communities.

Analyses of vOTU macro- and microdiversity revealed low genomic variation and strong purifying selection shaping viral populations in cold seep ecosystems. Macrodiversity, assessed by alpha-diversity indices (Shannon and Chao1), showed that seep type and sediment depth significantly structured viral communities (Kruskal-Wallis tests, *P* < 0.05), with sediment samples exhibiting markedly higher diversity than bottom-water (Shannon: 6.89 vs 5.38; Chao1: 6,425 vs 992; **Fig. S4a-b and Table S3**). Based on 20 samples (P1-P20) from site W07, Spearman correlation analyses revealed that Shannon diversity was not significantly correlated with any measured environmental variables. In contrast, Chao1 richness showed significant positive correlations with the concentration of potassium (K^+^; *P* < 0.05; **Fig. S4c)**. Furthermore, the overall vOTU community composition was significantly associated with several measured environmental variables, particularly sediment depth and ammonium (NH_4_^+^; Mantel’s *R* > 0.6 and *P* < 0.01; **Fig. 1d**). Cold seep viral populations exhibited low genome-level microdiversity: nucleotide diversity (π) ranged from 0 to 6.60 × 10^-3^ (mean: 2.62 × 10^-4^), and average SNP density was only 0.71 SNPs per kb (**Fig. 1e and Fig. S5a**). At the gene level, cold seep viral genes exhibited strong purifying selection, with most genes showing pN/pS < 0.05 (**Fig. 1f**) and strongly negative Tajima’s D values (often < -2.5; **Fig. S5b**). Together, these results indicate that cold seep viral populations experience strong evolutionary constraints and are shaped predominantly by purifying selection^33,34^.

### Extensive unclassified viral diversity with a substantial archaeal viral component

Taxonomic classification assigned 76.69% of GCSV vOTUs to a lineage. However, only 0.52% could be resolved to the order or family level **(Fig. 2a, Fig. S6 and Table S4**), highlighting that cold seep viruses remain highly undercharacterized and largely unclassified at finer taxonomic ranks. A total of 76.23% of vOTUs were assigned to the class *Caudoviricetes*, a dominant viral group in cold seeps^15,16^ and other ecosystems^4–6,30,35^, although most could not be resolved further. Among the vOTUs that were classified, *Zobellviridae* (n = 309), *Schitoviridae* (n = 212), and *Demerecviridae* (n = 97) were the most abundant families (**Fig. 2a and Table S4**). A small fraction (0.28%) of vOTUs, with an average genome length of 19.8 kb, were identified as nucleocytoplasmic large DNA viruses (NCLDVs) within *Nucleocytoviricota*. These viruses likely infect eukaryotic microorganisms such as protists and fungi^36^, suggesting potential ecological roles in cold seep eukaryotic communities.

**Figure 2.**
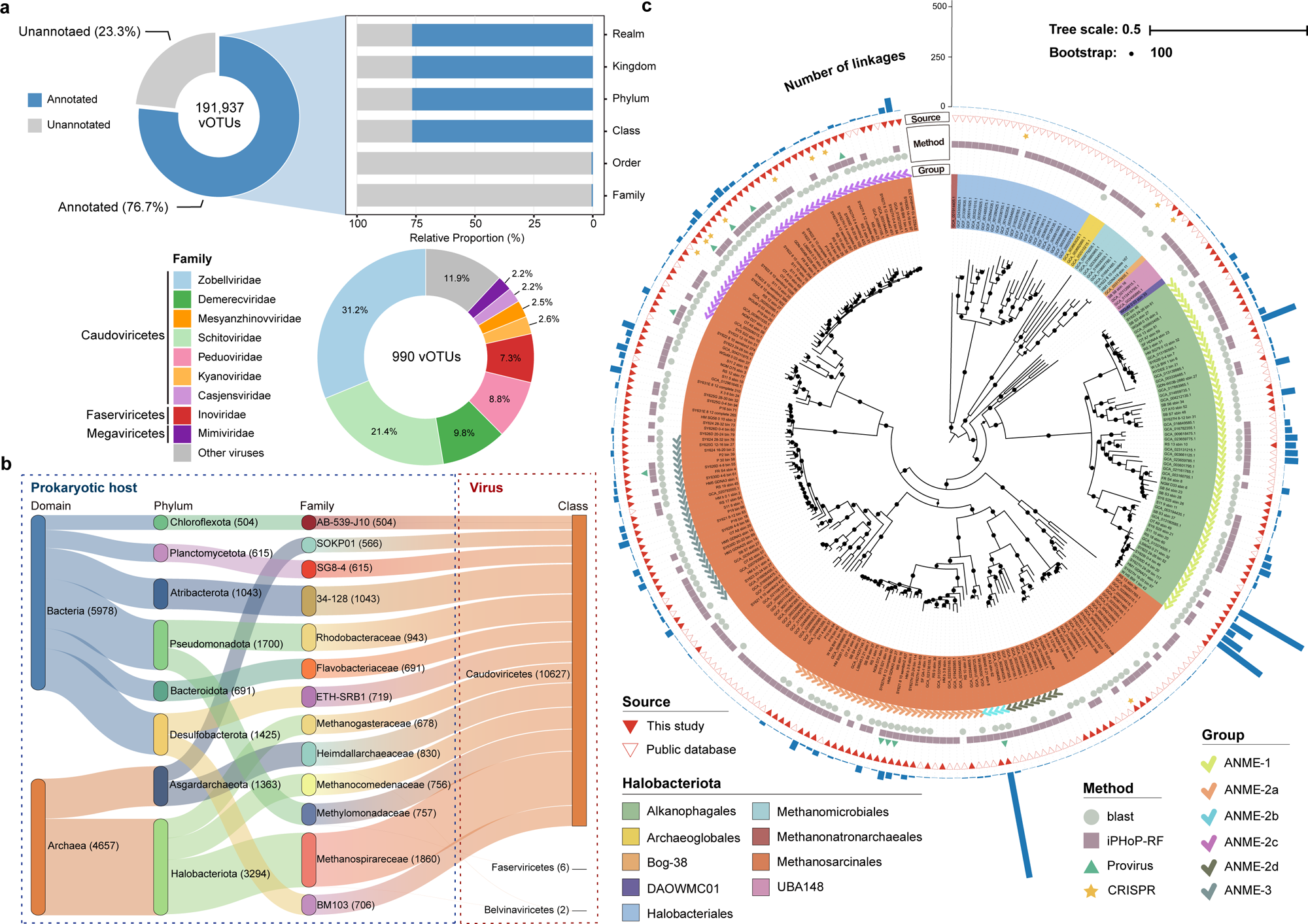
Taxonomic assignments and predicted hosts of vOTUs in the GCSV. **(a)** Taxonomic annotation of vOTUs. The upper pie chart summarizes the proportions of classified and unclassified vOTUs. The stacked bar plot displays the proportion of annotated (blue) and unannotated (grey) vOTUs across taxonomic ranks **(Table S4)**. The lower pie chart illustrates the family-level composition of classified vOTUs. Only vOTUs with family-level assignments were included, and the nine most abundant families are shown (color), with all remaining families grouped as “Others” (grey). **(b)** Sankey diagram illustrating predicted virus-host linkages. The left panel shows the major prokaryotic host groups (top eight phyla and top fifteen families), and the right panel shows the linked viral taxa at the class level. **(c)** Diversity of *Halobacteriota* hosts and associated viruses. Maximum-likelihood phylogenomic tree of *Halobacteriota* MAGs, constructed from a concatenated alignment of 53 archaeal single-copy marker genes. Background colors indicate taxonomic orders. From inside to outside, the tree rings show: (i) ANME subgroup classification; (ii) method of host-linkage inference; (iii) genome source; and (iv) number of vOTUs linked to each host genome. The scale bar indicates the number of substitutions per site and the node size is shown only for nodes with bootstrap support of 100.

Host prediction assigned putative hosts to 40,810 vOTUs (21.26%; **Table S5**). In total, 9,514 microbial genomes were linked to at least one virus, including 4,121 of the 7,010 genomes recovered from cold seep samples. Bacteria were the dominant predicted hosts (**Fig. 2b and Fig. S7**), spanning 123 phyla, with *Pseudomonadota* (7,130 linkages), *Desulfobacterota* (5,187), and *Bacteroidota* (4,214) being the most common (**Table S5**). Archaeal virus-host linkages accounted for 21.11% of all linkages, involving 8,557 vOTUs (4.46%) and 16 archaeal phyla. Among archaea, *Halobacteriota* were the most frequently predicted hosts (4,293 linkages; **Fig. 2c**), consistent with their dominance in cold seep microbiomes. Phylogenomic analyses revealed that *Halobacteriota* hosts included 172 anaerobic methanotrophic archaea (ANME) genomes (**Fig. S8a**). The GCSV contained 3,645 ANME-linked viruses, representing 89.14% of all *Halobacteriota*-associated viruses (**Fig. 2c and Table S5**). Viruses predicted to infect ANME spanned six subgroups, with ANME-1 being the most abundant (n = 1,913), followed by ANME-2b (n = 547) and ANME-2c (n = 516). Because anaerobic methane oxidation by ANMEs in cold seeps is primarily carried out in syntrophic partnerships with sulfate-reducing bacteria (SRB), we also examined the latter and identified 55 SRB genomes from three subgroups linked to 775 vOTUs (**Fig. S8b and Table S5**). These findings suggest that cold seep viruses may be associated with key microbial processes in methane oxidation and sulfate reduction.

Viruses predicted to infect *Asgardarchaeota* were also prevalent (n = 1,926), predominantly associated with *Sigynarchaeales*, *Heimdallarchaeales*, and *Thorarchaeales* (**Fig. S9**). In addition, we identified 412 and 617 viruses predicted to infect *Aenigmarchaeota* and *Nanoarchaeota*, respectively, both belonging to DPANN lineages characterized by ultra-small cells and reduced genomes that thrive in anoxic or extreme environments^37^. Machine-learning–based viral classification further indicated that potential archaeal viruses accounted for 7.60% of vOTUs at a MArVD2 probability ≥ 0.8 (**Fig. S3b; Table S2**), exceeding the proportion inferred from host linkages and suggesting substantial archaeal viral diversity yet to be taxonomically or ecologically resolved. Compared with other environments, e.g. soils^3^, groundwater^6^, and freshwater lakes^35^, cold seeps harbored a relatively high abundance of potential archaeal viruses, particularly lineages associated with *Halobacteriota* and *Asgardarchaeota*.

### Widespread uncharacterized proteins and ecologically adapted hallmark genes

From the 191,937 vOTUs in the GCSV, we predicted 4.82 million proteins, which were further clustered into 2.44 million non-redundant viral protein clusters (vPCs) at 50% sequence identity and 90% sequence overlap^38^. Representative sequences of each vPC were annotated using the protein language model ProstT5 ^39^ together with Foldseek^40^ searches against four structural databases, including Phold^41^, BFVD (The Big Fantastic Virus Database)^42^, PDB100 ^43^ and AlphaFold DB/Swiss-Prot v4 ^44^ (**Fig. 3a**). In total, 106,662 vPCs were assigned structural homologs across all databases. Functional categories derived from Phold annotations revealed that DNA/RNA/nucleotide metabolism (28.63%), head and packaging functions (21.87%), and tail proteins (20.26%) dominated the annotated space. Despite confident structural matches (E-value ≤ 10^-10^), the median sequence identity to known structures was only 24% (**Fig. 3b**), highlighting how structural conservation persists even when sequence similarity is no longer detectable. To complement structure-based annotation, vPCs were also searched against the eggNOG 5.0 database^45^ (**Fig. S10**) and enVhogDB^46^. Combining sequence-and structure-based approaches, 827,399 vPCs received at least one annotation. However, 66.1% remained uncharacterized **(Fig. 3c)**, underscoring the genetic novelty of cold seep viral communities^24^.

**Figure 3.**
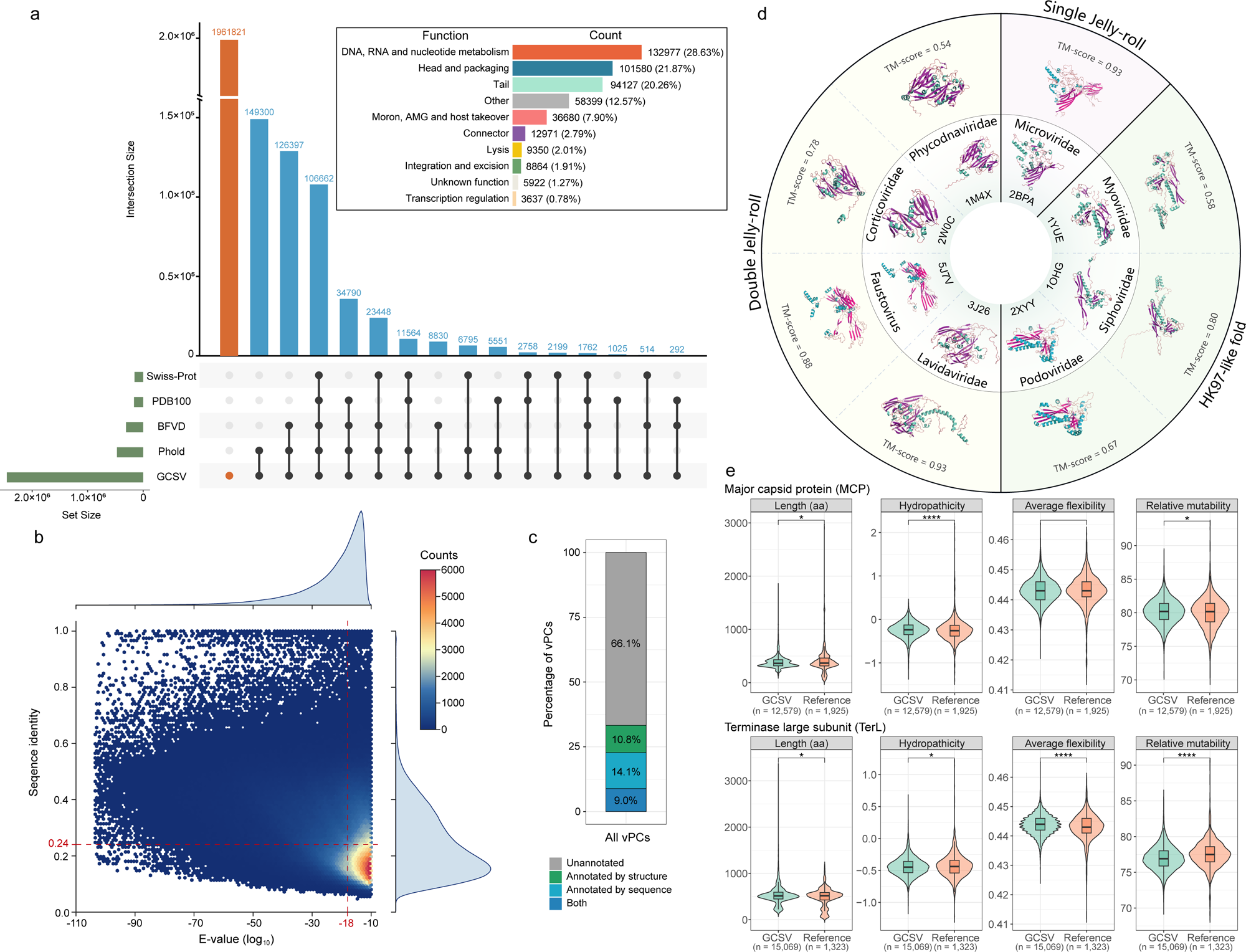
Structure-guided protein annotation and hallmark gene features in the GCSV. **(a)** Viral protein clusters were searched with Foldseek against PDB100, AFDB-Swiss-Prot, BFVD and Phold, retaining hits with E-value ≤ 10^-10^; the UpSet plot shows intersection sizes and per-set totals, and the inset summarizes functional categories of structurally annotated proteins. **(b)** Sequence identity versus log_10_(E-value) for best Foldseek matches across the four databases; red dashed lines indicate the medians (identity = 0.24, log_10_(E-value) = -18); marginal density plots show distributions. **(c)** Proportion of vPCs annotated using sequence-based and structure-based approaches. **(d)** Three canonical MCP folds—SJR, DJR and HK97-like. The inner ring shows PDB references; the outer ring shows GCSV MCP models superposed on references with TM-scores. **(e)** Violin plots compare MCP and TerL properties between GCSV (green) and reference proteins (orange), including length, hydropathicity, average flexibility and relative mutability. *n* indicates the number of proteins in each group. Group differences were tested using the Wilcoxon rank-sum test (*, *P* < 0.05; **, *P* < 0.01; ***, *P* < 0.001; ****, *P* < 0.0001).

Among viral hallmark proteins, we identified 12,579 major capsid proteins (MCPs) spanning a wide length range (119-1,827 aa; **Table S6**). Based on the classification framework of Krupovic et al.^47^ and PDB-derived structural annotations, 3,369 representative MCPs were confidently assigned to three canonical folds: single jelly-roll (SJR), double jelly-roll (DJR), and HK97-like (**Fig. 3d and Fig. S11a**). The majority of MCPs (n = 3,086) adopted the HK97-like fold typical of tailed dsDNA viruses (realm *Duplodnaviria*, class *Caudoviricetes*)^47,48^, consistent with their ecological dominance in cold seeps. DJR MCPs (n = 280) were affiliated with families such as *Mimiviridae*, *Phycodnaviridae*, and *Corticoviridae* within *Varidnaviria*^49^. By contrast, SJR MCPs were extremely rare (n = 3), likely reflecting limitations of standard metagenomic protocols in recovering ssDNA viruses^50^. Multimeric structure predictions further supported that HK97-like hexons and DJR triplexes formed stable capsomers (pTM = 0.75-0.88; **Fig. S11b**), supporting their functional relevance in cold seep viruses.

The terminase large subunit (TerL), another hallmark gene, was represented by 15,069 vPCs, with protein lengths ranging from 110 to 3,330 aa (**Table S7**). These proteins exhibited the canonical bipartite organization comprising an N-terminal ATPase domain and a C-terminal nuclease domain, with conserved ATP-binding motifs and nuclease catalytic residues (**Fig. S12**). Further structural comparisons revealed that structural conservation was markedly higher in domain-specific alignments (TM-score ≥ 0.77), which encompass the conserved residues, compared to full-length alignments (TM-score = 0.52-0.57). These results indicate that functional cores of cold seep viral TerLs remain stable, while the overall protein structures exhibit substantial diversity.

Although MCPs and TerLs retained conserved structural folds and catalytic features, both proteins exhibited distinct shifts in physicochemical properties and amino acid composition compared with reference proteins from BFVD^42^, Swiss-Prot^51^, and PHROG (Prokaryotic Virus Remote Homologous Groups)^52^ **(Fig. S13 and Table S8**). Both proteins, particularly MCPs, were significantly more hydrophilic (*P* < 0.05; **Fig. 3e**). This shift was driven by an enrichment of acidic residues (e.g., aspartic and glutamic acids), which facilitates the formation of stable hydration shells to protect structural integrity against high hydrostatic pressure and low temperatures^53^. Specifically, MCPs displayed significant glycine enrichment and proline depletion (*P* < 1×10^-5^; **Fig. S14**), thereby enhancing localized plasticity to counteract cold-induced rigidity^54^. Furthermore, TerLs exhibited significantly lower relative mutability compared with references (*P* = 2.71× 10^-33^; **Fig. 3e**), indicating strong purifying selection to remain conserved under extreme conditions. More broadly, significant divergence was observed across a total of 15 amino acid types for MCPs and 13 for TerLs (*P* < 0.05; **Fig. S14 and Fig. S15**). These shifts are consistent with possible environmental filtering or adaptation of viral hallmark proteins to cold seep conditions, while their deeply conserved structural frameworks are maintained.

### A functionally diverse repertoire of putative auxiliary metabolic genes

Using standard filtering criteria (see **Methods and materials**), we identified a diverse repertoire of 5,946 putative AMGs in the GCSV, representing 405 distinct functional annotations and encoded by 4,497 vOTUs (**Table S9**). At the KEGG level-2 tier (**Table S10**), these genes were mainly associated with metabolism of cofactors and vitamins (37.98%), glycan (27.82%), carbohydrate (13.08%), energy (12.80%), and amino acid (11.27%). These patterns indicate that cold seep viruses encode a broad repertoire of genes linked to diverse microbial metabolic processes.

We detected fifty putative AMGs associated with methane metabolism, including *pmoC*, which encodes a subunit of particulate methane monooxygenase (**Fig. 4**). Additional genes involved in formaldehyde oxidation and C1 assimilation (e.g., *fae, serA, psp, glyA*; **Table S9**) were also identified. Beyond methane, cold seep viruses carried putative AMGs involved in non-methane hydrocarbon degradation, including *alkB* and *cyp153* for aerobic alkane oxidation, as well as *ahyA* and *assA* for anaerobic alkane activation^55^. Cold seep viruses also encoded putative AMGs associated with central carbon metabolism, such as transketolase (*tkt*) in the Calvin-Benson cycle, pyruvate phosphate dikinase (*ppdK*) in the reductive TCA pathway, methylenetetrahydrofolate dehydrogenase (*folD*), and ribose-5-phosphate isomerase (*rpiAB*), suggesting possible links to central carbon metabolism. As cold seeps are chemosynthesis-driven ecosystems rich in chemotrophic microbes^56^, these genes may represent routes by which viruses could interact with host-associated primary production and carbon metabolism.

**Figure 4.**
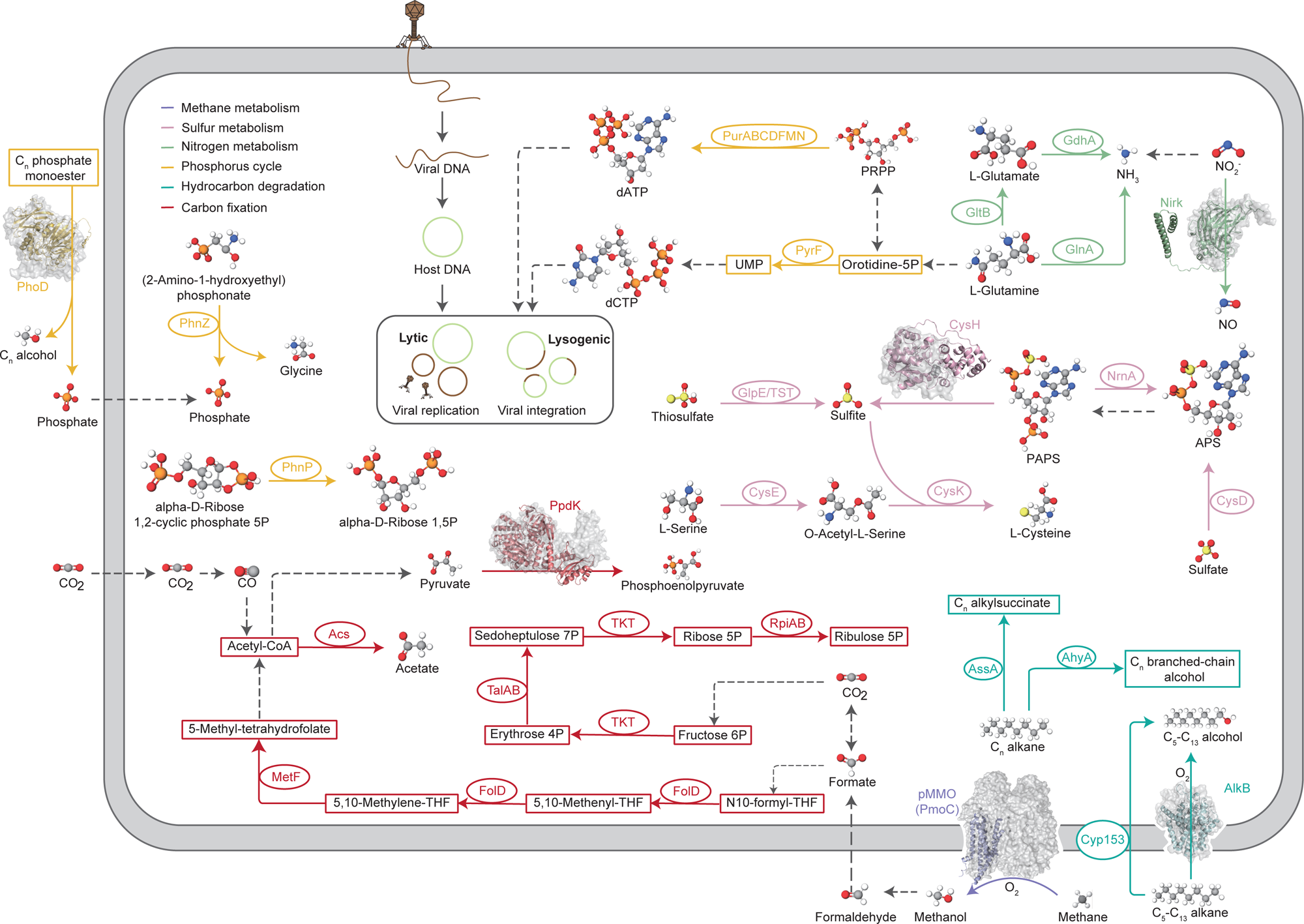
Metabolic potential encoded by cold seep viruses. Genes detected in the GCSV are colored according to their respective metabolic pathways, whereas undetected genes or pathway steps are shown in grey and indicated by dashed arrows. Genomic neighborhood analyses of representative displayed genes showed viral-gene flanking (**Fig. S16**), supporting their viral origin. Ovals represent AMGs (details in **Table S9**), and boxes denote the associated metabolites. For the protein structures shown, colored models correspond to AMG-encoded proteins identified in the GCSV, whereas grey models represent reference structures retrieved from the PDB. Details of structural alignments are provided in **Fig. S17**.

Cold seep viruses further carried putative AMGs implicated in multiple elemental cycles. In sulfur metabolism, *cysH* (PAPS reductase) was by far the most abundant putative AMG (n = 627), detected in viruses linked to hosts spanning 31 bacterial and archaeal phyla (**Table S5**). Additional assimilatory genes (e.g., *cysDEK*) and sulfurtransferases (TST) were also identified (**Fig. 4**), suggesting potential viral links to cysteine biosynthesis and thiol homeostasis. In nitrogen metabolism, we detected a *nirK* gene (copper nitrite reductase) associated with a *Methyloprofundus* host (**Fig. 4**), consistent with the known genomic content of this methanotrophic lineage^57^ and possible host-to-virus gene transfer. We also recovered genes involved in ammonium, nitrate, and nitrite assimilation or transformation (**Table S9**), including glutamine synthetase (*glnA*), glutamate synthase (*gltB*), asparagine synthase (*asnB*) and asparaginase (*ansB*). In phosphorus metabolism, we identified *phoD*, *phnP* and *phnZ*, which mediate organic phosphoester hydrolysis and phosphonate degradation, respectively (**Fig. 4**). These genes were primarily linked to *Desulfobacterota*, including SRB (**Table S5**). Cold seep viruses also encoded genes involved in nucleotide biosynthesis, including genes from purine metabolism (*guaA, purABCDFMN*) and pyrimidine metabolism (*dcd, dut, nrdADE, pyrEF, thyA, tmk*).

### A broad viral anti-defense arsenal against multi-layered prokaryotic immunity

To overcome the wide variety of prokaryotic defense systems present in cold seep environments^58^, viruses in the GCSV encode an extensive repertoire of anti-defense mechanisms. Across the dataset, 4,944 vPCs were predicted to encode anti-defense genes spanning 13 distinct systems (**Fig. 5a and Table S11**). These vPCs were derived from 4,571 vOTUs, with 85.6% assigned to *Caudoviricetes* and only a small fraction to *Varidnaviria* (**Fig. 5b**). Among all anti-defense systems, anti-CRISPR (23.9%), anti-restriction-modification (anti-RM; 21.7%), and anti-toxin-antitoxin (anti-TA; 12.4%) were most abundant, followed by anti-CBASS, anti-Thoeris, and NADP systems. Within the anti-CRISPR category, the *vcrx091-093* operon (n = 1,064) was dominant. This module mediates recombination-based repair of CRISPR-induced double-strand breaks, promoting CRISPR-Cas evasion^59^. The anti-RM system was largely driven by *dmt* (n = 294), whereas *mga47* (n = 458) accounted for the majority of anti-TA genes. Other common antagonists included *adnd_p0020* (anti-Dnd; n = 380), *ntase* (anti-CBASS; n = 338), and *tad2* (anti-Thoeris; n = 264). Anti-defense elements were unevenly distributed across viral genomes. We identified 319 vOTUs carrying multiple anti-defense genes, with the most enriched genome encoding four distinct systems comprising five anti-defense genes (**Fig. 5c**). The most frequent combinations were anti-CRISPR + anti-RM and anti-CRISPR + anti-CBASS.

**Figure 5.**
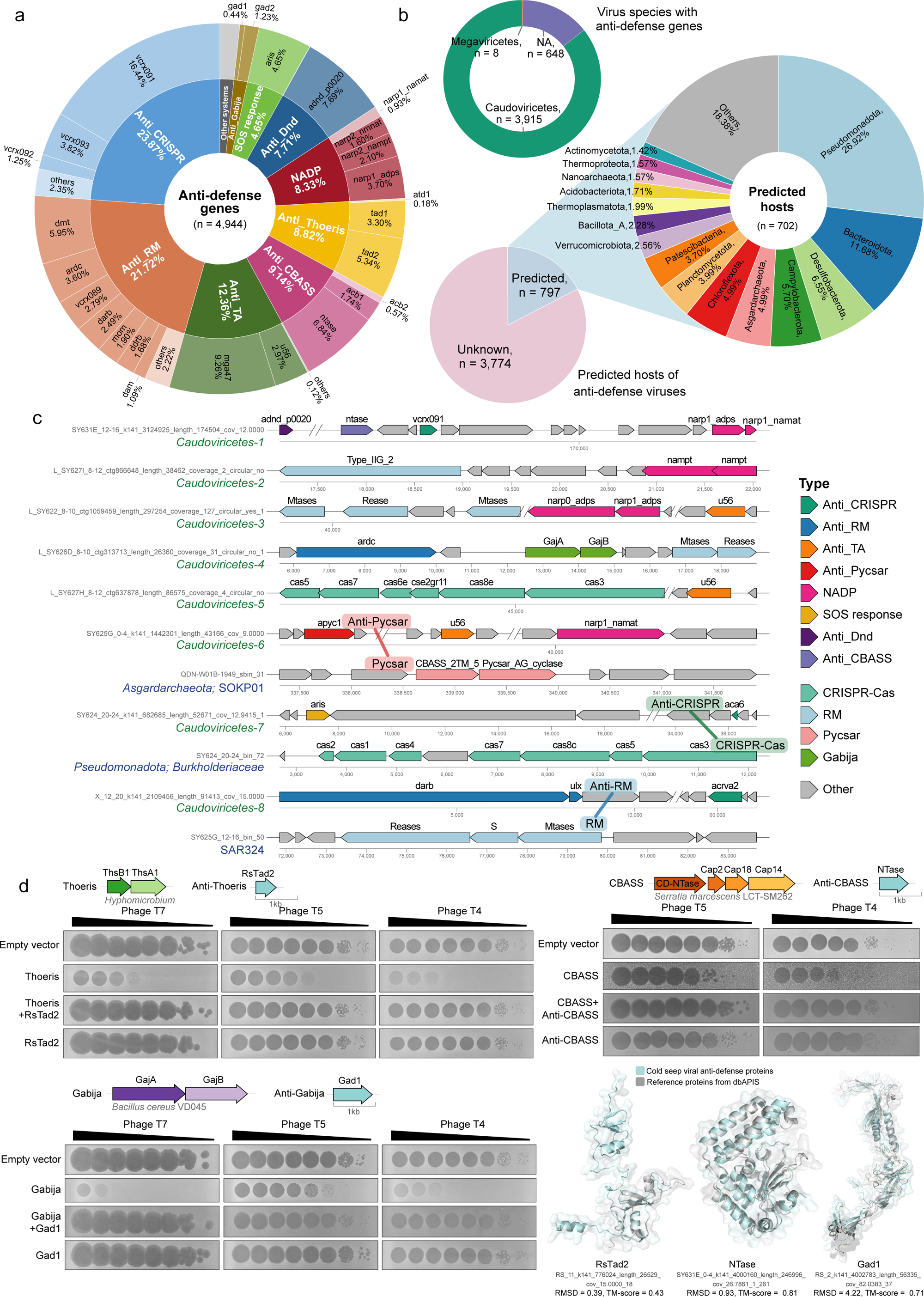
Anti-defense systems encoded by cold seep viruses. **(a)** Distribution of 4,944 anti-defense vPCs across 13 systems in the GCSV. Outer rings highlight representative genes (e.g., *vcrx091-093*, *dmt*, *tad2*, *gad1*, NTase). **(b)** Taxonomic composition of anti-defense viruses (class level) and predicted hosts (phylum level). **(c)** Representative genomic loci illustrating co-localization of anti-defense and host defense genes, including multi-system co-occurrence and examples consistent with predicted virus-host pairs. **(d)** Functional validation of selected anti-defense candidates. Plaque assays of *E. coli* B expressing defense systems, anti-defense genes, or both, challenged with phages T7, T5, and T4 using the small-drop method. Defense systems include Thoeris (ThsB1-ThsA1) from *Hyphomicrobium* sp., Gabija (GajAB) from *Bacillus cereus* VD045, and type-I CBASS from *Serratia marcescens*. Experiments were performed in triplicate with similar outcomes. Bottom right: AlphaFold3 structural models of RsTad2, Gad1, and NTase aligned with dbAPIS reference proteins.

Virus-host linkage analysis revealed 873 virus-host pairs involving anti-defense–encoding viruses. These pairs corresponded to 797 vOTUs (17.4% of all anti-defense–carrying viruses) linked to 702 host MAGs spanning 61 bacterial and archaeal phyla (**Fig. 5b**). The predicted hosts were dominated by *Pseudomonadota* (26.9%), followed by *Bacteroidota* (11.7%), *Desulfobacterota* (6.6%), *Campylobacterota* (5.7%), *Asgardarchaeota* (5.0%), and *Chloroflexota* (5.0%). Host genomes collectively encoded 4,523 defense genes representing 161 systems, primarily RM and CRISPR-Cas (**Table S12**). Among these, 138 virus-host pairs showed direct correspondence between viral anti-defense genes and host defense systems, most commonly anti-RM vs RM (n = 109) and anti-CRISPR vs Cas (n = 23). For example, one *Caudoviricetes* genome encoded an anti-Pycsar protein Apyc1 alongside the anti-TA and NADP proteins, while its predicted *Asgardarchaeota* host carried a complete Pycsar system (**Fig. 5c**). Defense genes were also detected within viral genomes themselves (**Fig. S18 and Table S12**). Notably, 60 vOTUs encoded both defense and anti-defense systems. These systems rarely targeted the same immunity pathways (**Fig. S19 and Table S12**). For instance, the representative *Caudoviricetes* genomes encoded the anti-TA gene *u56*, as well as CRISPR-Cas or RM systems (**Fig. 5c**). This pattern suggests that these viruses may not only suppress host immunity but also deploy their own defense modules to compete against other mobile genetic elements^60^.

To validate the functional activity of anti-defense genes, we selected one representative from each of three abundant systems, including anti-Thoeris (RsTad2), anti-Gabija (Gad1), and anti-CBASS (NTase), and expressed them in *E. coli* B either alone or co-expressed with their respective defense systems (Thoeris ThsB1-ThsA1 from *Hyphomicrobium* sp.; Gabija GajAB from *Bacillus cereus* VD045; type I CBASS from *Serratia marcescens*). In all cases, cells expressing defense systems alone showed strong protection (severe plaque reduction), whereas co-expression with the matched anti-defense gene restored plaque formation, demonstrating efficient defense neutralization (**Fig. 5d**). Specifically, RsTad2 and Gad1 abolished Thoeris- and Gabija-mediated immunity against all three phages tested, respectively, and NTase suppressed CBASS-mediated resistance to phages T5 and T4. Mechanistically, Tad2 sequesters Thoeris immune signaling molecules^61^, Gad1 forms octameric assemblies that block Gabija-mediated DNA cleavage^61^, and NTase produces competing cyclic dinucleotides that inhibit CBASS effector activation^62^. As expected, independent expression of these anti-defense genes did not alter phage resistance to the host compared to the empty vector control, confirming that their effect on bacterial immunity is strictly dependent on the presence of a cognate defense system. Structural models predicted by AlphaFold3 showed strong similarity to known anti-defense proteins, providing further support for accurate functional annotation (**Fig. 5d, bottom right**).

### Diverse viral lysins with potential for antimicrobial applications

Lysins are essential viral effectors that degrade bacterial cell walls during progeny release^63^ and typically consist of an enzymatically active domain (EAD) and a cell wall binding domain (CBD)^64,65^. From the GCSV, we identified 15,688 lysins from vPCs using structural alignment and DeepMineLys^66^. Among these, 5,338 carried high-confidence lysin domains and were more likely to exhibit antibacterial activity (**Table S13**). A total of 187 vOTUs encoded multiple lysins, with some containing up to five distinct EAD types. Additionally, 10,350 lysins lacked recognizable domains (**Fig. 6a**). While some of these sequences may contain domains not represented in current reference databases^65,67^, it is also likely that a portion of these represents prediction artifacts of automated tools. Compared with existing viral lysin datasets, which largely derive from human microbiomes or general environmental viromes^65–70^, the GCSV represents one of the most extensive deep-sea viral lysin resources to date.

**Figure 6.**
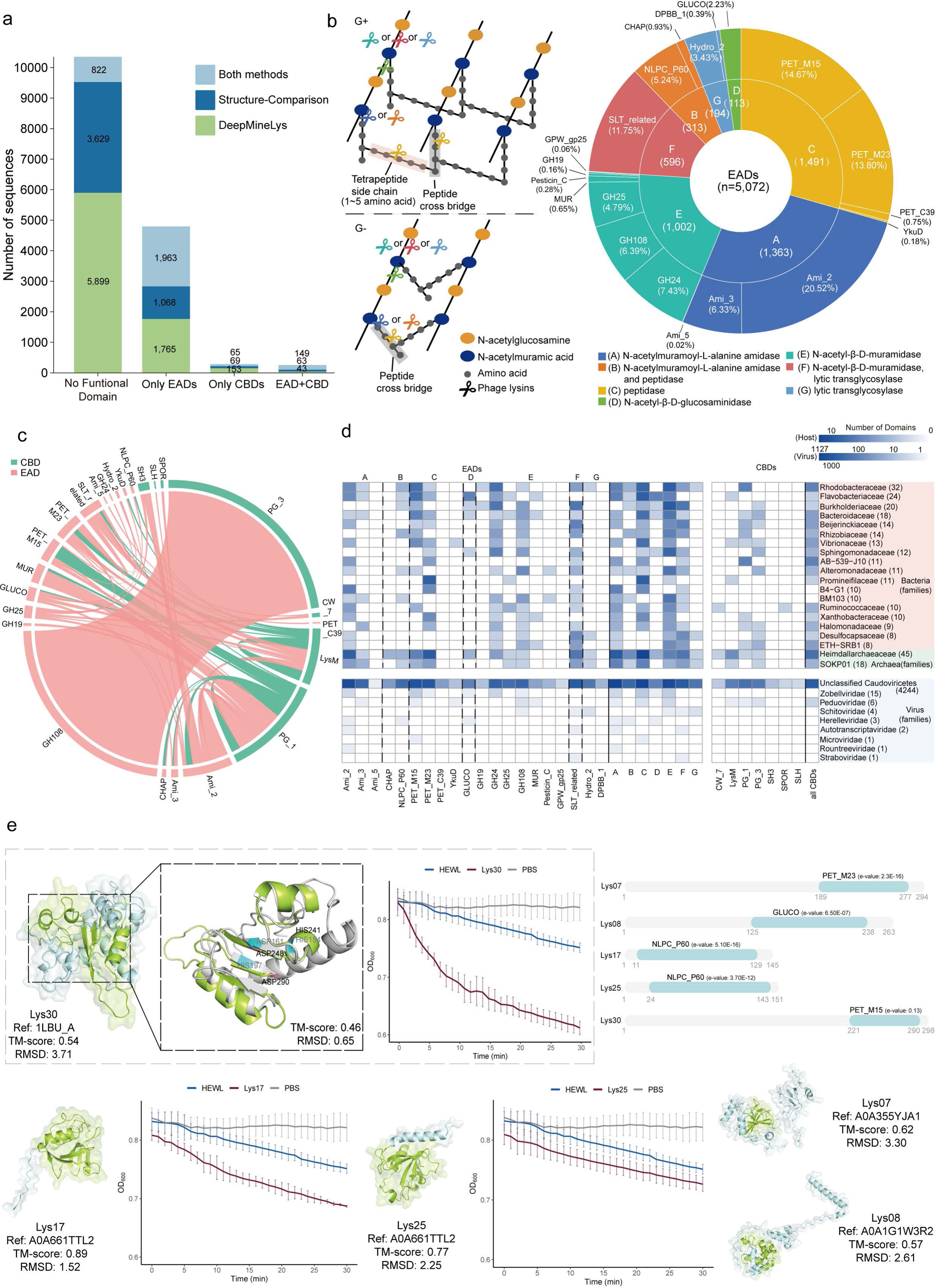
Diversity, architecture, and host distribution of lysins encoded by cold seep viruses. **(a)** Classification of lysins based on domain composition. Different bar colors indicate annotation approaches used to identify these lysins. EADs, enzymatically active domains; CBDs, cell wall binding domains. **(b)** Cleavage positions of major EAD enzymatic classes (left) and classification of the 5,072 EADs into seven activity groups comprising 20 domain families (right). **(c)** Domain adjacency patterns in lysins. Pink links denote EADs and green links denote CBDs, with link thickness reflecting co-occurrence frequency. **(d)** Distribution of EAD and CBD domain families across predicted viral host families (top 20 shown) and viral families (bottom). Heatmap intensity reflects log_10_-transformed domain counts, with sequence numbers indicated in parentheses. **(e)** Experimental validation of five lysins. Domain architectures are shown at top right (EADs in light blue). Predicted AlphaFold3 structures highlight EADs (light green) and remaining regions (sky blue), with closest structural homologs and TM-score/RMSD values noted. Lytic activity against *Bacillus subtilis* ATCC 6051 is shown for three representative lysins, with HEWL and PBS as positive and negative controls. Values represent mean ± SD from three independent experiments. Structural alignments of the Lys30 EADs with reference proteins (grey) are presented, with catalytic residues labeled in black for lysins and grey for reference proteins. Additional lysin activity curves are provided in **Fig. S23**.

Domain-level analyses revealed substantial catalytic diversity and domain combinations among cold seep viral lysins (**Fig. 6a and Fig. S20**). Most sequences encoded at least one EAD (n = 4,775), followed by CBD-only sequences (n = 276) and lysins containing both an EAD and a CBD (n = 217). EADs encompassed 20 domain families grouped into seven catalytic classes^71^ (**Fig. 6b and Table S14**), dominated by amidases (Ami_2, Ami_3), peptidases (PET_M15, PET_M23), lytic transglycosylases (SLT-related), and muramidases (GH24). CBDs exhibited lower diversity, representing seven families enriched in PG_1 (42%), PG_3 (28%), and LysM (26%) (**Fig. S21**). Domain adjacency analysis revealed multiple pairings, with GH108 + PG_3 in EAD-CBD being the most frequent (**Fig. 6c**). Given that peptidoglycan composition varies across microbial taxa, lysins with multiple catalytic or binding modules may facilitate broader host specificity and enhanced lytic efficiency^71^.

Taxonomic mapping uncovered uneven distributions of lysin domains **(Fig. 6d and Fig. S22)**. Among viruses, Ami_2 showed the broadest distribution across viral families, whereas some EADs (e.g., GLUCO, Hydro_2) were restricted to one or two families. CBDs showed more restricted distributions. Across predicted hosts, half of the top 20 families were associated with five or more EADs. Viruses predicted to infect *Heimdallarchaeaceae, Rhodobacteraceae* and SOKP01 encoded relatively diverse lysin repertoires, covering nearly all catalytic classes. Among these, *Heimdallarchaeaceae*-associated viruses showed the highest domain diversity, including 12 EADs across seven catalytic types. By contrast, viruses linked to B4-G1, AB-539-J10, and *Promineifilaceae* encoded only a few domains. Several EADs, including Ami_2, PET_M23, and SLT-related, were widespread across > 25 host phyla, whereas some (e.g., Ami_5, PET_C39) were restricted to *Bacillota*_A and *Patescibacteria*, respectively. Similarly, CBDs such as LysM and PG_1 exhibited broad host ranges across 20 phyla, while CW_7, SLH and SPOR were restricted to *Bacillota*_A and *Asgardarchaeota*. These observations potentially indicate that some domains are more broadly distributed, whereas others are more specific. However, these patterns may partly reflect methodological and reference biases, as domains with higher overall frequency also tended to show broader distributions, and some domains may remain uncharacterized.

To evaluate antimicrobial potential, thirty lysin candidates were selected for experimental testing. Five (Lys07, Lys08, Lys17, Lys25, Lys30) were successfully expressed in soluble form in *Escherichia coli*, as confirmed by SDS-PAGE, whereas the remaining candidates displayed poor solubility. All five purified lysins exhibited detectable hydrolytic activity against *Bacillus subtilis* ATCC 6051 and four deep-sea-derived bacterial strains (*Bacillus tequilensis* GS003, *Bacillus altitudinis* GS004, *Mesobacillus thioparans* GS011, and *Bacillus tequilensis* GS013), as evidenced by time-dependent reductions in the optical density (OD) of bacterial cultures. Lys07, Lys08, and Lys25 exhibited activities comparable to the benchmark hen egg white lysozyme (HEWL), whereas Lys17 and Lys30 showed substantially higher activity **(Fig. 6e and Fig. S23a-b**). Colony-counting assays confirmed strong antibacterial effects for all five lysins, with inhibition rates of 50-90%, while Lys17 and Lys30 showed activities comparable to or exceeding that of HEWL (**Fig. S23c**). The correspondence between hydrolytic and bactericidal effects indicates that peptidoglycan degradation is the primary killing mechanism. Given that all these lysins are of deep-sea origin, their lytic activity was further validated using a panel of deep-sea-derived Gram-positive bacterial strains. In *Bacillus tequilensis* GS003 (**Fig. S24a**), the lytic activities of Lys07 and Lys08 were substantially higher than that of HEWL (positive control), while the activity of Lys25 was higher in *Mesobacillus thioparans* GS011 (**Fig. S24c**). These lysins also displayed lytic activity against *Bacillus altitudinis* GS004 and *Bacillus tequilensis* GS013 (**Fig. S24b and S24d**), although the activity was weak. Further structural analyses revealed that the EAD of Lys30 (PET_M15) diverged from known reference proteins, as reflected by low structural alignment scores (TM-score < 0.6; **Fig. 6e**) and weak sequence-based domain assignments (E-values > 0.01; **Table S15**). Despite retaining predicted catalytic residues, Lys30 featured more loops and fewer α-helices relative to its homologs. Such structural flexibility may enhance enzymatic performance under low-temperature and high-pressure conditions^72^, although this remains to be experimentally validated.

## Discussion

The GCSV, a catalog comprising 191,937 species-level vOTUs and 2.44 million vPCs, represents a > 66-fold increase in vOTU counts compared to the previous cold-seep viral dataset (2,885 vOTUs)^15^ and stands as one of the largest deep-sea viral resources to date. Analysis of this dataset reveals a viral ecosystem that is taxonomically distinct, evolutionarily constrained, functionally diverse, and shaped by complex virus-host conflicts. Fewer than 0.3% of GCSV vOTUs overlap with public viral datasets from seven diverse ecosystems^4–6,29–32^, highlighting cold seeps as major reservoirs of undescribed viral lineages. This pattern is further underscored by the substantial fraction of unclassified *Caudoviricetes*, indicating that many cold seep viral lineages remain poorly represented in current taxonomic frameworks. In addition, the diverse predicted archaeal viruses, spanning *Halobacteriota*, *Asgardarchaeota*, and multiple DPANN lineages, provide a resource for future investigations of archaeal virus diversity and virus-archaea interactions in methane-rich deep-sea environments.

Overall, only 21.26% of the vOTUs in the GCSV could be confidently assigned to a host. This relatively low host-assignment rate highlights a current limitation of *in silico* host prediction for deep-sea viromes, likely due in part to the limited representation of uncultivated deep-sea-derived microbial genomes in the reference database. Furthermore, given the ecological prominence of archaea in these habitats^13,73^, their associated viruses are expected to be well represented. However, the proportion of inferred archaeal virus-host associations was substantially lower than that of bacterial associations. This pattern may reflect methodological biases rather than the true abundance of archaeal viral infections in cold seep ecosystems. Consequently, it is probable that a portion of the unassigned viral sequences within the GCSV represents previously uncharacterized archaeal viruses.

Despite this high genomic novelty, cold seep viruses exhibit remarkably low microdiversity and strong purifying selection. This indicates that their evolution is not shaped by rapid mutation-based diversification as observed in surface oceans^4^, but is relatively constrained by long-term genomic stability likely driven by low temperature, high pressure, and chemically stratified sediments^16,74^. While viral micro-diversity remains highly constrained, viral macro-diversity and overall community structure may be influenced by local environmental gradients. As demonstrated by analysis at site W07, viral community composition and richness are significantly related to multiple geochemical factors. However, these inferences remain limited because comparable *in situ* geochemical parameters are unavailable for most sites. Future comprehensive studies incorporating broader and higher-resolution environmental metadata are required to further elucidate the complex virus-environment interactions in cold seep ecosystems.

The extensive fraction of unannotated viral proteins identified in the GCSV, with nearly two-thirds of protein clusters lacking structural or sequence-based annotation, highlights substantial unresolved viral genetic diversity in cold seep systems and the potential for discovering new molecular mechanisms. The GCSV also provides an integrated view of viral functions associated with cold seep biogeochemistry. Hallmark proteins such as MCPs and TerLs retain conserved structural folds while showing distinct physicochemical patterns relative to reference datasets, consistent with possible adaptation to extreme environments^53^.

Beyond these genetic features, this study also provides insights into potential connections between viruses and cold seep metabolic processes. The GCSV contains a diverse repertoire of putative AMGs compared to other deep-sea viral datasets^75^, including genes associated with carbon fixation, methane oxidation, hydrocarbon degradation under both oxic and anoxic conditions, and multiple elemental cycles. In this context, the nitrogen-related signal at site W07 illustrates a possible link between viral metabolic potential and local geochemical conditions: viral community composition was associated with sediment NH_4_ ^+^ levels, and the GCSV contained several nitrogen-related genes, including *glnA*, which encodes glutamine synthetase involved in ammonium assimilation^76^. This association remains tentative owing to the limited geochemical coverage and the lack of direct functional validation.

Cold seep viruses also encode a diverse repertoire of anti-defense genes. Predicted matches between viral anti-defense modules and host defense systems, together with experimental validation, support lineage-specific antagonism and ongoing virus-host arms races^77^. These functionally active immunity inhibitors highlight cold seep viruses as a rich genetic reservoir for developing new molecular tools and potentially novel therapeutic strategies^78,79^. The discovery of lysins with previously unrecognized structural domains further emphasizes the biotechnological value of cold seep viruses. The diversity of catalytic units, modular architectures, and cold-adapted structural features suggests robust lytic capabilities. Unlike conventional antibiotics, lysins hydrolyze peptidoglycan directly, reducing the likelihood of resistance, making cold seep lysins promising candidates for next-generation antimicrobial development and broader applications in environmental and industrial biotechnology^68^.

Collectively, the GCSV provides an expansive and foundational resource for ecological, evolutionary, and biotechnological research. By revealing extensive viral novelty, putative functional genes, and potential signatures of adaptation to extreme environmental conditions, this catalog expands the boundaries of the known virosphere and establishes cold seeps as previously overlooked centers of viral innovation and potential ecological influence. Moving forward, integrating this resource with multi-omics analyses, experimental validation, and functional characterization will be essential for translating these genomic insights into practical applications and for clarifying the roles of viruses in deep-sea chemosynthetic ecosystems.

## Methods and materials

### Sample collection, DNA extraction, and sequencing

In November 2023, we collected 79 sediment samples and two bottom water samples from ten cold seep sites in the South China Sea, with a maximum water depth of 1,830.1 m. Sediment samples were obtained using short push cores deployed by the deep-sea manned submersible *Shen Hai Yong Shi*, with one core collected at each site. The sediment cores reached depths of up to 34 centimeters below seafloor (cmbsf) and were sectioned at 4-cm intervals, except for the core from site SY630, which was sectioned at 2-cm intervals. At two sites (SY626 and SY628), the *in situ* microbial filtration and fixation (ISMIFF) apparatus^80^ was deployed directly above mussel beds using the submersible. The device was equipped with 0.22-µm, 142-mm polycarbonate filters and operated for approximately 4-6 hours to collect *in situ* bottom water samples. Detailed information on sampling locations and sediment intervals is provided in **Table S1**. All samples were stored at -80°C until DNA extraction.

For each sediment sample, ∼5 g of material was used for DNA extraction, and each filter membrane was cut into small pieces prior to extraction. DNA was extracted using the DNeasy PowerSoil Pro Kit (47014, QIAGEN) following the manufacturer’s instructions. To obtain longer DNA fragments suitable for long-read sequencing, the cell lysis step was optimized by adding 60 µL Buffer SC (TIANGEN BIOTECH, China). Metagenomic sequencing was conducted on the DNBSEQ-T7 high-throughput sequencing platform for 74 samples (72 sediment and two bottom water samples), generating a total of 3.64 Tb of short-read data. Additionally, seven representative sediment samples were selected for long-read sequencing based on DNA yield. These samples were sequenced using the PacBio Revio platform for HiFi long-read sequencing, yielding a total of 0.5 Tb of PacBio reads.

### Metagenomic dataset collection and integration

A total of 191 cold seep sediment metagenomic datasets were collected from our previous publications^14,81^. In addition, six sediment metagenomes from the Hikurangi Margin methane seeps (HYDEE) were downloaded from the National Center for Biotechnology Information Sequence Read Archive (NCBI-SRA). We also included 36 bottom water metagenomes collected from the Haima cold seep^82–84^. By integrating the newly obtained samples described above with previously published metagenomic datasets, the GCSV dataset comprises 276 sediment samples and 38 bottom water samples originating from 18 geographically diverse cold seeps. These cold seeps include (**Fig. 1a**): Eastern North Pacific (ENP), Santa Monica Mounds (SMM), Western Gulf of Mexico (WGM), Eastern Gulf of Mexico (EGM), Northwestern Gulf of Mexico (NGM), Scotian Basin (SB), Haakon Mosby mud volcano (HM), Mediterranean Sea (MS), Laptev Sea (LS), Shenhu area (SH), Haiyang4 (HY4), Qiongdongnan Basin (QDN), Xisha Trough (XST), Haima cold seep (HM1, HM3, HM5, HM_SQ, S11, SY5, SY6, W_HM_1, W_HM_2, W_HM_3), Site F cold seep (RS, SF, FR, SF_SQ), Okinawa Trough (OT), SY region (SY621, SY622, SY623, SY624, SY625, SY627, SY628, SY630, SY631, W07, L_SY), and HYDEE. These cold seeps encompass multiple geological types, including oil and gas seeps, methane seeps, gas hydrate systems, asphalt volcanoes, and mud volcanoes. The maximum water depth among these sites was 3,408 m, and the deepest sediment sample was collected at 68.55 meters below seafloor (mbsf). Detailed information on sampling sites, sediment depths, sequencing metadata and data sources is available in **Table S1**. Additionally, the geochemical parameters for site W07 were obtained from our previous study^81^.

### Metagenomic quality control and assembly of short- and long-read data

Quality control of the newly generated short-read data was performed using fastp^85^ (v0.23.4; default parameters) to obtain high-quality clean reads. Clean reads from each metagenome were individually assembled using MEGAHIT^86^ (v1.1.3; default parameters) and contigs ≥ 1,000 bp were retained for downstream binning analyses. For previously published datasets, quality control and assembly followed the procedures described in our earlier study^14^ (**Table S1**). Briefly, either the Read_QC module within the metaWRAP^87^ pipeline or fastp was used for quality filtering, and clean reads were assembled using MEGAHIT with the same parameters. In total, 97.5 million contigs (≥ 1,000 bp) were obtained. Long-read (PacBio HiFi) data were assembled using metaMDBG^88^ (v1.0; default parameters), producing 2.4 million contigs longer than 1,000 bp.

### Metagenomic binning and non-redundant MAG catalog construction

Binning of the newly generated short-read assemblies was performed using the binning module of metaWRAP^87^ (v1.3.2; parameters: -metabat2, -maxbin2, -concoct, - universal), along with the single_easy_bin mode of SemiBin2 ^89^ (v2.1.0; default parameters). For previously published datasets, binning was conducted using metaWRAP, SemiBin (v1.4.0), and Rosella^90^ (v0.4.1; default parameters), as detailed in **Table S1**. For each assembly, bins generated by all binning tools were integrated and refined using the Bin_refinement module within metaWRAP (parameters: -c 50 -x 10). For PacBio long-read assemblies, binning was performed with pb-metagenomics-tools using the HiFi-MAG-Pipeline^91^ (v3.2.0, pb-metagenomics-tools).

The completeness and contamination of all refined bins were evaluated using CheckM^92^ (v1.2.1). Applying thresholds of ≥ 50% completeness and ≤ 10% contamination, a total of 17,981 metagenome-assembled genomes (MAGs) were recovered from 314 samples. All MAGs were dereplicated at the species level using dRep^93^ (v2.5.4) with a 95% average nucleotide identity (ANI) cutoff, resulting in 7,010 representative MAGs. Taxonomic classification of these MAGs was performed using GTDB-Tk^94^ (v2.4.0) against the Genome Taxonomy Database (release 220)^95^.

### Identification and clustering of viral genomes

Viral genomes were identified from 6.2 million contigs (≥ 5,000 bp) obtained from short-read and long-read assemblies, including two million contigs from PacBio HiFi assemblies, using a two-step workflow (**Fig. S1**). In the first step, contigs were screened with three viral identification pipelines: geNomad (v1.8.0)^96^, VirSorter2 (v2.2.4)^97^, and VIBRANT (v1.2.1)^98^. Candidate viral genomes were retained following previously published criteria^3,29^. (1) For geNomad predictions, contigs of 5-10 kb were required to have a virus score ≥ 0.9, at least one viral hallmark gene, and a virus-marker enrichment > 2.0, whereas contigs ≥ 10 kb were required to have a virus score ≥ 0.8 and at least one hallmark gene or a virus-marker enrichment > 5.0. (2) High-confidence VirSorter2 predictions (score ≥ 0.8 and ≥ 1 hallmark gene) were retained. (3) Contigs co-identified by VirSorter2 (score ≥ 0.5) and VIBRANT were further screened by CheckV^99^ (v1.0.1); contigs with viral_gene = 0 and checkv_quality = “Not-determined” were removed. Putative proviruses identified at this stage were retained as candidate viral sequences for subsequent refinement rather than being directly accepted as proviral genomes.

In the second step, CheckV was used to refine the candidate viral contigs. Contigs were retained if viral_gene > 0. For contigs in which viral_gene = 0, retention required satisfying at least one of the following criteria^100^: (i) host_gene = 0; (ii) VirSorter2 score ≥ 0.95; (iii) more than two VirSorter2 hallmark genes; (iv) geNomad score ≥ 0.95; (v) more than two geNomad hallmark genes. After these filtering steps, the “viruses.fna” and “proviruses.fna” files generated by CheckV were merged, resulting in 313,811 viral sequences. These sequences were then clustered into non-redundant viral (nr-viral) sequences at 95% average nucleotide identity (ANI) and 85% alignment fraction (AF) of the shorter genome using CheckV scripts, yielding 193,465 nr-viral sequences for subsequent refinement.

To further reduce false positives, identify proviruses, and remove host contamination, the nr-viral contigs were subjected to additional curation. Plasmid sequences were identified using PLASMe^101^ (v1.1; -m high-precision), and 54 detected plasmids were removed. Concatemers were identified following the metaVR pipeline^102^ and its script (https://pages.jgi.doe.gov/imgvr5-399e0f/pages/additional_analysis.html), resulting in 1,416 sequences, which were excluded as likely assembly artifacts.

The dataset was then divided into “virus” and “provirus” subsets. The virus set included sequences classified as viral rather than proviral by geNomad, VirSorter2, and CheckV (n = 179,711). The provirus set comprised: (i) sequences classified as viruses by geNomad or VirSorter2 but as proviruses by CheckV (n = 6,322), and (ii) sequences classified as proviruses by geNomad or VirSorter2 (n = 5,888). The latter were re-evaluated following Dahlman et al.^103^, whereby proviral regions predicted by geNomad and VirSorter2 were first extracted, then processed with the CheckV contamination module to remove microbial regions. Overlapping host-derived regions were subsequently merged using IRanges^104^ (v2.28.0).

Both groups of proviral sequences were further processed following the metaVR pipeline^102^ approach to remove regions containing complete rRNA genes, identified by barrnap^105^ (v0.9; parameters: --kingdom bac, arc, euk; --evalue 1e-10). After trimming all contaminating regions, sequences shorter than 1 kb were discarded, resulting in 12,420 proviruses. These were clustered at 95% ANI and 85% AF, yielding 12,226 non-redundant proviruses. For the six sequences exceeding 500 kb that remained after the refined filtering process, we performed manual curation by inspecting and visualizing their gene-level annotations based on CheckV’s results **(Figure S25)**. These sequences were retained in the final dataset as they exhibited relatively more viral genes than host genes. Finally, the virus set and nr-provirus set were merged, resulting in a final dataset of 191,937 vOTUs.

### Comparison with other viral databases based on vOTUs

To assess the novelty and cross-biome distribution of cold seep vOTUs, all representative vOTU genomes were compared against seven publicly available viral sequence datasets, including mudflat intertidal sediments^29^, soils^30^, groundwater^6^, GOV 2.0 ^4^, the Arctic Ocean surface microlayer^31^, deep subsurface oil reservoirs^5^, and the rumen microbiome^32^. Cold seep vOTUs were searched against each database using BLASTn^106^. For each BLASTn alignment, ANI and alignment fraction (AF) were calculated based on pairwise nucleotide identity and aligned length relative to the shorter sequence. A vOTU was considered present in a given dataset if the best hit satisfied both thresholds: ≥ 95% ANI and ≥ 85% AF.

### Viral taxonomy, abundance, and diversity profiling

Taxonomic classification of viral genomes was performed using three complementary approaches, including geNomad^96^ (v1.8.0), vConTACT3 ^107^ (v3.0.0b74; default parameters) and VITAP^108^ (v1.5; default parameters). The geNomad classifications followed the International Committee on Taxonomy of Viruses (ICTV) Taxonomy Release MSL39, while vConTACT3 was run using database v223 ^109^ derived from NCBI Virus RefSeq release 223, and VITAP was run using ICTV Taxonomy Release MSL40.v1. To ensure consistency across tools, all classifications were harmonized to ICTV Release MSL40.v1 ^110^.

Taxonomic assignments were determined using a majority consensus approach among three tools. At each taxonomic rank, a lineage was only retained if supported by at least two tools, and assignments lacking such consensus were designated as unclassified. To further enhance the accuracy of taxonomic assignments, we refined the results by integrating structure-based MCP annotations (detailed below). Specifically, for vOTUs harboring proteins confirmed as DJR-MCPs through structural alignment against PDB references (TM-score ≥ 0.7; **Figure S26**), but initially classified as *Caudoviricetes*, the taxonomic assignment was revised accordingly to *Varidnaviria* to maintain consistency between structural and taxonomic features.

The abundance of vOTUs in each sample was calculated with the contig mode of CoverM^111^ (v0.6.1; parameters: --trim-min 0.1 --trim-max 0.90 --min-read-percent-identity 0.95 --min-read-aligned-percent 0.75). For diversity analyses, clean reads were mapped to vOTU sequences using Bowtie2 ^112^ to generate BAM files, and MetaPop^113^ (v0.0.42; default parameters) was applied to estimate both macrodiversity and microdiversity of viral populations. Microdiversity was assessed at the contig level, using metrics such as nucleotide diversity (π) and the number of single nucleotide polymorphisms (SNPs), and at the gene level, using metrics including Tajima’s D and the pN/pS ratio.

### Host prediction for prokaryotic viruses

In accordance with the Virus-Host Database^114^, vOTUs belonging to non-prokaryotic lineages, including *Herpesvirales*, *Cirlivirales*, *Nucleocytoviricota*, and *Virophaviricetes*, were excluded from the host prediction pipeline. Host prediction for vOTUs was performed using the iPHoP^115^ pipeline (v1.3.3; default parameters), which integrates multiple host prediction strategies^116–119^. Prior to prediction, a customized reference database was constructed by incorporating 7,010 MAGs recovered from cold seep samples into the iPHoP database (iPHoP_db_Aug23_rw) using the “add_to_db” function of iPHoP, and all MAG taxonomies were standardized to GTDB release r220. To reduce false-positive assignments, we applied a hierarchical filtering scheme based on iPHoP results. For proviruses, the microbial genome containing the integrated viral region was directly assigned as the host. When multiple host predictions were obtained for a vOTU, they were ranked in the following order of priority^35^: (1) provirus located within a host genome, (2) CRISPR-based iPHoP prediction, (3) BLAST-based iPHoP prediction, and (4) iPHoP-RF. Within each tier, if multiple candidate hosts were present, the prediction with the highest iPHoP confidence score was selected as the final host assignment. Additionally, the proportion of archaeal viruses in the GCSV was estimated using the machine learning-based tool MArVD2 ^120^ (v0.11.9; default parameters), with probability ≥ 0.8 used as the high-confidence threshold for archaeal virus classification.

To resolve the phylogenetic placement of viral hosts, we constructed four separate phylogenomic trees using GTDB-Tk based on concatenated sets of archaeal or bacterial single-copy marker genes: *Halobacteriota*, *Asgardarchaeota*, ANME (ANME-1, ANME-2a/2b/2c/2d, ANME-3), and SRB (Seep-SRB1a, Seep-SRB1g, Seep-SRB2). All trees were visualized and annotated in iTOL^121^.

### Structure- and sequence-based viral protein annotation

Viral proteins encoded by vOTUs were predicted using Prodigal^122^ (v2.6.3; -p meta), and the resulting protein sequences were clustered into viral protein clusters (vPCs) using MMseqs2 ^123^ easy-cluster module (v13.45111; parameters: --min-seq-id 0.5 -c 0.9 --cov-mode 1) at 50% sequence identity and 90% coverage^38^. The representative sequence from each vPC was used for downstream analyses. Protein annotation was primarily based on structure-guided functional prediction, in which amino acid sequences were converted into 3Di-tokens using the ProstT5 protein language model^39^ and aligned against structural databases with Foldseek^40^. Two complementary strategies were applied. First, Phold^41^ (v0.2.0), which uses the ProstT5 + Foldseek workflow, was employed to search against a curated database of more than 1.36 million predicted viral protein structures with high-quality functional labels. Second, the ProstT5 + Foldseek workflow (Foldseek release 10; parameters: --gpu 1 -c 0.5 --cov-mode 0) was used to search three major structural databases: BFVD^42^, PDB100 ^43^ and AlphaFold DB/Swiss-Prot v4 ^44^. For each query sequence, the annotation with the lowest E-value in each database was retained as the top hit, and only hits with E-value ≤ 1×10^-10^ were considered for further analysis.

Two sequence-based approaches were additionally used to annotate viral proteins. First, EggNOG-mapper^124^ (v2.1.9; parameters: -m diamond --tax_scope Viruses) was used to search against the eggNOG 5.0 database^45^. Second, MMseqs2 search^123^ (v13.45111; parameters: --start-sens 2 -s 7 --sens-steps 3 -a) was applied to search the enVhog database^46^, retaining only matches with ≥ 30% identity and ≥ 80% coverage. For each protein, the best annotation was selected by prioritizing the lowest E-value and highest sequence identity.

### Prediction and alignment of protein structures

Protein structures, including monomeric proteins and protein complexes, were predicted using AlphaFold3 ^125^ (https://alphafoldserver.com/). Pairwise structural alignments were performed with TM-align^126^ (v20170708; default parameters). All protein structures were visualized using PyMOL^127^ (v3.0.0).

### Physicochemical properties of MCPs and TerLs

Viral hallmark proteins, including MCPs and TerLs, were identified from structure-based gene annotation results. Reference proteins were collected from PHROG, BFVD and Swiss-Prot databases. Physicochemical properties of each protein, including hydrophobicity, polarity, relative mutability, transmembrane tendency, and refractivity, were calculated using the ProtScale tool (https://web.expasy.org/protscale/)^128^. Amino acid composition was calculated as the frequency of each residue relative to the total number of amino acids in the protein sequence.

### Auxiliary metabolic gene identification

Based on geNomad and VirSorter2 annotations, vOTUs containing viral hallmark genes were selected for putative AMG identification. Candidate AMGs were predicted using two complementary approaches, DRAM-v^129^ (v1.3.5) and VIBRANT, followed by multiple genomic-context and functional filters to remove genes located within host-derived regions and generate a curated catalog of putative AMGs. For DRAM-v–based identification, vOTU sequences were first processed with VirSorter2 (parameter: --prep-for-dramv) to generate the required input files, and then annotated using DRAM-v (default parameters). The resulting annotations were curated following the criteria of Tian et al^17^. Genes flanked on both sides by viral-like or viral hallmark genes (auxiliary scores ≤ 3) and carrying the DRAM-v metabolism flag (M) were considered permissive candidates. These candidates were further filtered by removing those derived from viral genomes lacking both tRNA regions (detected with tRNAscan-SE^130^ v1.23; parameters: -G) and inverted or direct repeats adjacent to the genome ends (predicted with EMBOSS einverted^131^ v6.6.0). We also removed candidates located on viral contigs that contained mobile genetic elements or genes that could facilitate nonspecific acquisition of microbial metabolic genes, including transposons, lipopolysaccharide island–associated genes (glycosyltransferases, nucleotidyl transferases, carbohydrate kinases, and nucleotide-sugar epimerases), endonucleases, integrases, and plasmid stability genes. These elements were identified based on the results of DRAM-v.

Candidate AMGs predicted by VIBRANT were curated following the criteria of Zhou et al^35^: (1) AMGs located at contig edges were removed; (2) AMGs with KEGG or Pfam v-scores ≥ 1 were excluded; (3) AMGs whose two upstream or downstream flanking genes all had KEGG v-scores < 0.25 were removed; and (4) genes annotated with COG categories T or B were excluded. The remaining VIBRANT-derived AMGs were retained as curated putative AMG candidates. Hydrocarbon-degrading genes were further identified using hmmsearch^132^ (parameter: --cut_nc) against the CANT-HYD database^55^ and BLASTp^133^ (parameter: -evalue 1e-10) against HMDB^134^. All candidate hydrocarbon-degrading genes were manually verified to ensure that each gene was flanked by viral genes, as annotated by Phold, confirming their viral genomic context.

Candidates passing the relevant DRAM-v, VIBRANT or hydrocarbon-gene filters were combined. Genes involved in queuosine biosynthesis (*queC*, *queD*, *queE* and *queF*) and DNA cytosine methylation (*dcm*) were subsequently excluded because these genes may participate in virus-centered processes, including viral genome modification, translation and defense evasion, rather than the auxiliary modulation of host metabolism^135^. The remaining candidates were compiled into the final curated catalog of putative AMGs.

### Anti-defense and defense gene identification

Viral anti-defense genes were predicted across all viral genes using two complementary approaches, and their corresponding vPC representative sequences were subsequently extracted: (1) predicting with AntiDefenseFinder^18^ (v2.0.2; parameter: --db-type gembase); and (2) homology searching against the APIS database^136^ using DIAMOND^137^ blastp (v2.0.8; parameter: -e 1e-10 --id 30) and HMMER^132^ (v3.3.2; parameter: -E 1e-10). To ensure high-confidence assignments, only matches to experimentally validated APIS genes were retained. For each query sequence, only the hit with the lowest E-value was retained from both DIAMOND blastp and HMMER results before merging, and conflicting annotations were resolved by prioritizing the HMMER assignment. Anti-defense system and gene names were standardized following the AntiDefenseFinder nomenclature. Consistent with previous studies^138^, all *acriia7* hits were reclassified as *tad2*. Defense genes present in viral and host genomes were identified using DefenseFinder^139^ (v2.0.2; parameter: --db-type gembase).

### Heterologous expression and activity verification of anti-defense genes

Coding sequences of representative anti-defense genes (RsTad2, Gad1, and a viral nucleotidyltransferase, NTase) and their cognate defense operons (Thoeris, Gabija, and CBASS) were synthesized and cloned by GenScript (Gibson Assembly) into pQE82L-derived expression vectors. Defense systems included the Thoeris operon (ThsB1-ThsA1) from *Hyphomicrobium* sp., the Gabija operon (GajAB) from *Bacillus cereus* VD045, and a type-I CBASS operon from *Serratia marcescens*. Constructs carrying (i) defense only, (ii) anti-defense only, (iii) defense together with the matched anti-defense gene, or (iv) an empty vector (negative control) were transformed into *Escherichia coli* B (ATCC® 11303™), and transformants were selected using appropriate antibiotics and DNA sequencing.

Phage-challenge assays were performed using the small-drop plaque assay^140,141^. Cultures were challenged with bacteriophages T7 (*Podoviridae*), T5 (*Siphoviridae*), and T4 (*Myoviridae*). Single colonies were inoculated into lysogeny broth (LB) supplemented with antibiotics and grown at 37°C to OD_600_ ≈ 0.6, after which protein expression was induced with 0.2 mM isopropyl β-D-1-thiogalactopyranoside (IPTG) for ∼1 h. For top agar overlays, 500 µL of the induced culture was mixed with 14.5 mL of LB soft agar (0.5%) containing antibiotics and 0.1 mM IPTG, and poured onto LB plates. Serial phage dilutions were prepared and spotted in 4-µL drops: T7 (10^-1^-10^-8^), T5 (10^0^-10^-7^), and T4 (10^0^-10^-7^). Plates were incubated at 25°C for ∼16-18 h and imaged. Phage-resistance phenotypes were scored by assessing plaque formation: reduction or absence of plaques indicated active defense, whereas plaque restoration upon co-expression of the cognate anti-defense gene indicated neutralization of the defense system. Anti-defense constructs alone were also tested to confirm the absence of intrinsic protective activity. All assays were performed in ≥ 3 independent biological replicates with similar results.

### Viral lysin identification and domain analysis

vPCs were searched against the PhaLP database^65^ (v2021_04; https://www.phalp.org) using MMseqs2 ^123^ (parameter: --min-seq-id 0.2), and sequences sharing ≥ 20% identity with reference lysins were retained. These sequences were further analyzed using DeepMineLys^66^, a convolutional neural network-based framework for phage lysin identification. Proteins annotated as endolysins or virion-associated lysins (VALs) by DeepMineLys were considered candidate lysins. In addition, proteins predicted as lysins by the Phold workflow or through Foldseek-BFVD structural matches were incorporated into the candidate lysin catalog. All candidate lysins were subsequently annotated using InterProScan^142^ (v5.0.39; parameters: -goterms -iprlookup -pa), which queries the InterPro database^143^, integrating signatures from 13 member databases. Only domain hits with E-value ≤ 1e-5 were retained. The identification and classification of these domains were performed following Criel et al.^65^ and Koposova et al.^67^ For proteins containing two or more functional domains, domain-linkage patterns were constructed and visualized as chord diagrams in R.

### Heterologous expression, purification and activity verification of lysins

The DNA sequences encoding 30 lysins (designated Lys01-Lys30) were codon-optimized for expression in *Escherichia coli* and individually cloned into the pET28a vector with an N-terminal His-tag using the *NdeI* and *XhoI* restriction sites (SynbioB, Tianjin, China). The codon optimization of the lysin genes was performed using the internal optimization software provided by SynbioB, following the codon usage preferences of *Escherichia coli*. Specifically, high-frequency codons selected from the Codon Usage Table **(Table S16)** were adopted for the optimization. The recombinant plasmids were transformed into *E. coli* BL21 (DE3) competent cells. Transformed cells were inoculated into LB medium supplemented with 50 mg/L kanamycin and incubated at 37°C with shaking at 220 rpm. When the optical density at 600 nm (OD_600_) reached 0.6-0.8, the cultures were immediately cooled on ice for 30 min. Protein expression was induced by the addition of IPTG to a final concentration of 0.1 mM, followed by incubation at 12°C for 24 h. Cells were harvested by centrifugation at 8,000 rpm for 10 min at 4°C, resuspended in lysis buffer (20 mM Tris–HCl, pH 7.5, 300 mM NaCl), and disrupted by ultrasonication. The lysate was centrifuged at 12,000 rpm for 1 h at 4°C, and the supernatant was loaded onto a Ni–NTA agarose column pre-equilibrated with lysis buffer. The column was washed sequentially with elution buffers containing 25 mM and 50 mM imidazole, and the target proteins were eluted with 150 mM imidazole in the same buffer. Eluted fractions were concentrated to 2.5 mL using an Amicon Ultra-15 centrifugal filter (10 kDa MWCO; Millipore) and dialyzed against PBS (pH 7.4) at 4°C for 12 h. Protein purity was assessed by 12% SDS–PAGE, and bands were visualized using Coomassie Brilliant Blue R-250 staining.

The peptidoglycan hydrolytic activity of purified lysins was evaluated using *Bacillus subtilis* ATCC 6051 and four deep-sea-derived bacterial strains (*Bacillus tequilensis* GS003, *Bacillus altitudinis* GS004, *Mesobacillus thioparans* GS011, and *Bacillus tequilensis* GS013) as the substrates following the protocol of Fu et al^66^. The deep-sea-derived strains were isolated from surface sediments (0-1 cm) of the North Pacific Abyssal Plain during the DY89 cruise (July 2025; multi-corers) using Marine Agar 2216E culture medium^144^. All strains were cultured in LB medium at 37°C for 5 h, diluted 50-fold into fresh LB, and grown to mid-logarithmic phase (OD_600_ = 0.5-0.6). Cells were pelleted, washed with PBS (pH 7.4), and resuspended to OD_600_ = 0.8-1.0. Each lysin was added to 150 µL of bacterial suspension at a final protein concentration of 0.1 mg/mL. The OD_600_ decrease was monitored continuously for 30 min using a Spark multimode microplate reader (Tecan, Austria). Commercial HEWL (Sangon Biotech, China) served as the positive control, and PBS was used as the negative control. All assays were performed in triplicate, and results are expressed as mean ± SD.

To further assess antibacterial activity, a colony-counting assay was performed. Briefly, 50 µL of each lysin (1 mg/mL) was mixed with 150 µL of *B. subtilis* ATCC 6051 suspension and incubated at 37°C for 1 h. Aliquots (50 µL) of each reaction mixture were plated on LB agar (1%) and incubated overnight at 37°C. Colony-forming units (CFU) were counted, and antibacterial activity was calculated using:

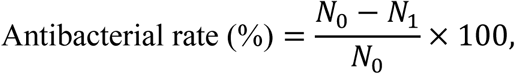

where *N_0_* and *N_1_* represent the CFU counts from the control and lysin-treated groups, respectively. All experiments were conducted in triplicate, and the values are reported as mean ± SD.

### Statistical analysis

Statistical comparisons of viral diversity parameters (macrodiversity and microdiversity) across habitats, seep types, and depth groups were performed using Kruskal-Wallis rank-sum tests, followed by two-sided pairwise Wilcoxon rank-sum tests with Benjamini-Hochberg correction for multiple comparisons, implemented in the rstatix R package^145^ (v0.7.0). Boxplots were generated using ggplot2 ^146^ (v3.3.6), where the center line represents the median, the bounds of the box represent the 25th and 75th percentiles (IQR), and the whiskers extend to 1.5 × IQR. Distinct letters above each box indicate statistically significant differences (*P* < 0.05).

The relationship between genome size and SNP abundance was calculated and visualized in R^147^ (v4.2.3). To minimize the influence of outlier contig lengths, contigs below the 1st percentile or above the 99th percentile of length distribution were excluded. From the remaining dataset, 90,000 contigs were randomly subsampled for visualization. Average SNP frequency was calculated as the total number of SNPs divided by the cumulative length of all retained contigs and expressed as SNPs per kilobase (SNPs/kb).

To explore the relationships between viral community composition and environmental parameters, Mantel’s tests were performed on 20 samples (P1-P20) collected from site W07 using the mantel_test() function in the linkET R package^148^ (v0.0.7.4), with significance assessed using 999 permutations. Pairwise Spearman correlations among environmental variables were calculated using the correlate() function and p-values were adjusted for multiple comparisons using the Benjamini-Hochberg FDR correction. The associations between viral alpha diversity indices (Shannon and Chao1) and individual environmental parameters were similarly evaluated using Spearman correlation with FDR correction. All analyses and visualizations were implemented in the linkET R package (v0.0.7.4).

Comparisons of physicochemical properties and amino acid composition between viral hallmark proteins in this study and reference proteins were visualized using violin plots overlaid with boxplots, and statistical significance was assessed using two-sided Wilcoxon rank-sum tests with Benjamini-Hochberg correction in R (v4.2.3).

## Data availability

Newly generated metagenomic sequencing data have been deposited under the European Nucleotide Archive (ENA) BioProject PRJEB10xxxx. Microbial MAGs and vOTU sequences generated in this study have been deposited under the ENA BioProject PRJEB11xxxx. Additional derived data, including vPCs and predicted AMGs generated in this study have been deposited on Figshare (https://doi.org/10.6084/m9.figshare.xxxxxxxx). This paper also analyzes previously published, publicly available metagenomic datasets. Detailed source information and accession identifiers for these datasets are provided in **Table S1**.

## Code availability

The code used in this study is available via GitHub.

## Acknowledgements

This work was supported by National Key R&D Program of China (No. 2024YFC2816200), National Natural Science Foundation of China (No. 42376115, No. 32470036, No. 32100025, and No. 42406109), Natural Science Foundation of Xiamen City (No. 3502Z202373076), Natural Science Foundation of Fujian Province (No. 2023J06042), Natural Science Foundation of Hubei Province (No. JCZRQNA202600209), Young Top-notch Talent Cultivation Program of Hubei Province, and Fundamental Research Funds for the Central Universities (No. 2662025SYPY003). We thank Lingdong Shi, Chuwen Zhang, Qiuyun Jiang, Chengpeng Li, and Wenqing Shi for valuable discussions and constructive suggestions on the manuscript. We are grateful to Jiwei Li and Haili Ran for assistance with sample collection and DNA extraction, and to Yuehong Wu, Hong Cheng, and Ling Cao for providing the deep-sea-derived bacterial strains. We also thank the National Key Laboratory of Agricultural Microbiology Core Facility for technical support with the EnVision microplate reader and protein purification, and Shaoran Zhang for assistance with instrument operation.

## Author contributions

X.D. and X.L. conceived and designed the study. X.L. performed the majority of metagenomic analyses, prepared figures, extracted DNA, and conducted sample preparation for sequencing. X.G. analyzed and identified viral lysins. L.X. and R.C. performed anti-defense gene assays and lysin functional validations. Z.L., J.L., J.W., L.L., and Y.C. contributed to parts of the metagenomic processing, data analysis, and figure preparation. Y.H. assisted with the collection, integration, and preprocessing of global metagenomic datasets, including the processing of long reads. Y.P. and F.W. provided scientific input and critical revisions. X.L., X.D., and X.G. wrote and revised the manuscript with contributions from all authors.

## Supplemental information

Supplementary figures: Figures S1-S26

Supplementary tables: Tables S1-S17

