## Supplementary Figures 1-26 for "A global atlas of deep-sea cold seep viruses uncovers extensive genomic novelty and biotechnological potential"

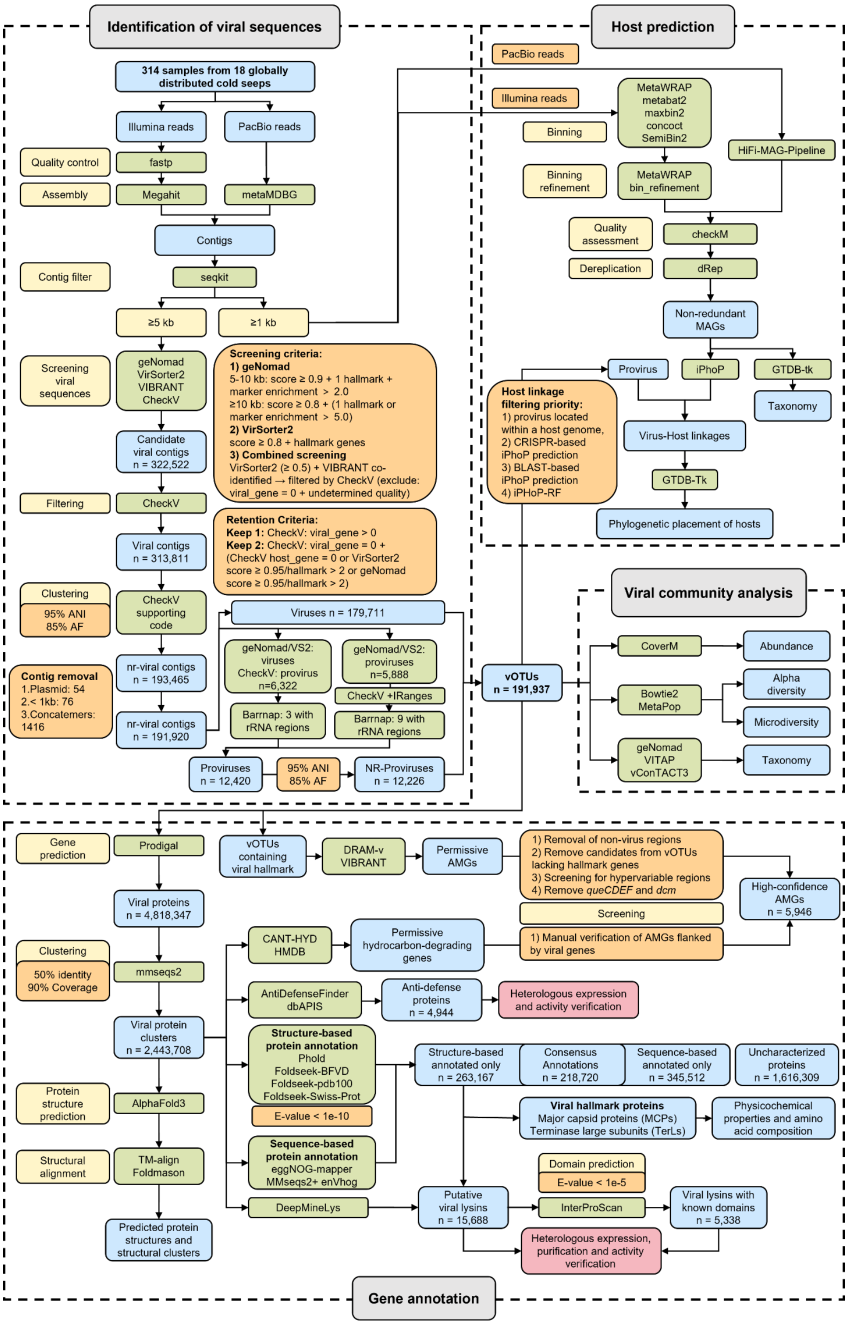


**Figure S1. Overview of the computational pipeline used to recover, classify, and functionally characterize viral genomes from cold seep metagenomes.** The workflow integrates four main steps: (1) identification of viral genomes through quality control, assembly, and screening with multiple tools; (2) viral host prediction based on cold seep metagenome-assembled genomes (MAGs) and reference genomes; (3) viral community analyses including abundance, diversity, and taxonomy; and (4) functional annotation of viral proteins, highlighting auxiliary metabolic genes, viral hallmark proteins, anti-defense proteins and viral lysins.

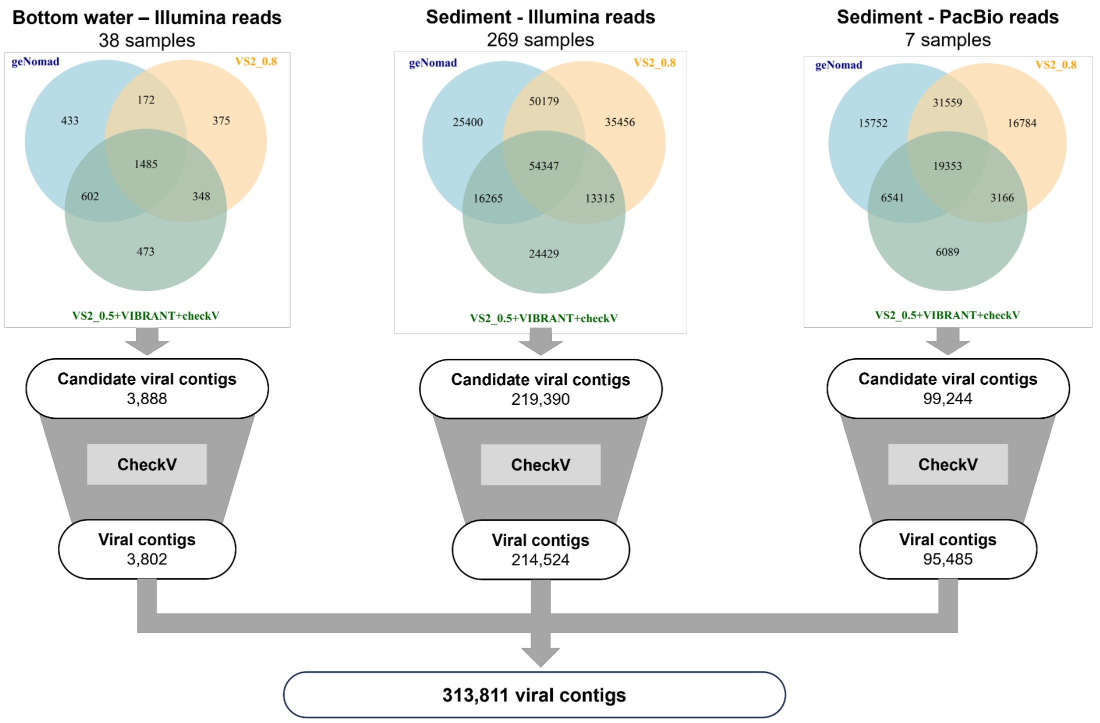


**Figure S2. Identification and quality assessment of viral contigs.** Candidate viral contigs were detected using geNomad, VirSorter2 (VS2), and VIBRANT, and subsequently filtered with CheckV for quality refinement (**Methods**). Venn diagrams illustrate the overlap among different detection tools, and funnel plots summarize the number of sequences retained after filtering. In total, 313,811 viral contigs were recovered from 314 metagenomes.

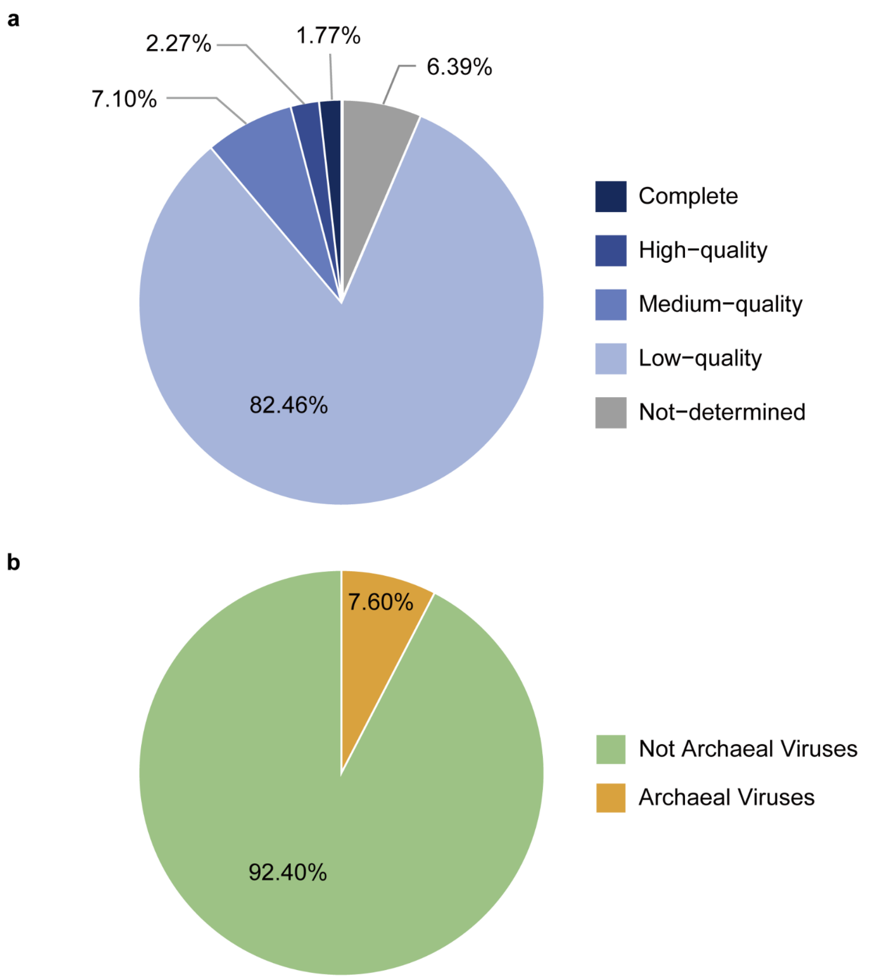


**Figure S3. Basic characteristics of the vOTU dataset from cold seep metagenomes.** **(a)** Distribution of vOTU completeness categories as assessed by CheckV. **(b)** Proportion of archaeal viruses identified by MArVD2 with a probability threshold of ≥ 0.8. Detailed data are provided in **Table S2**.

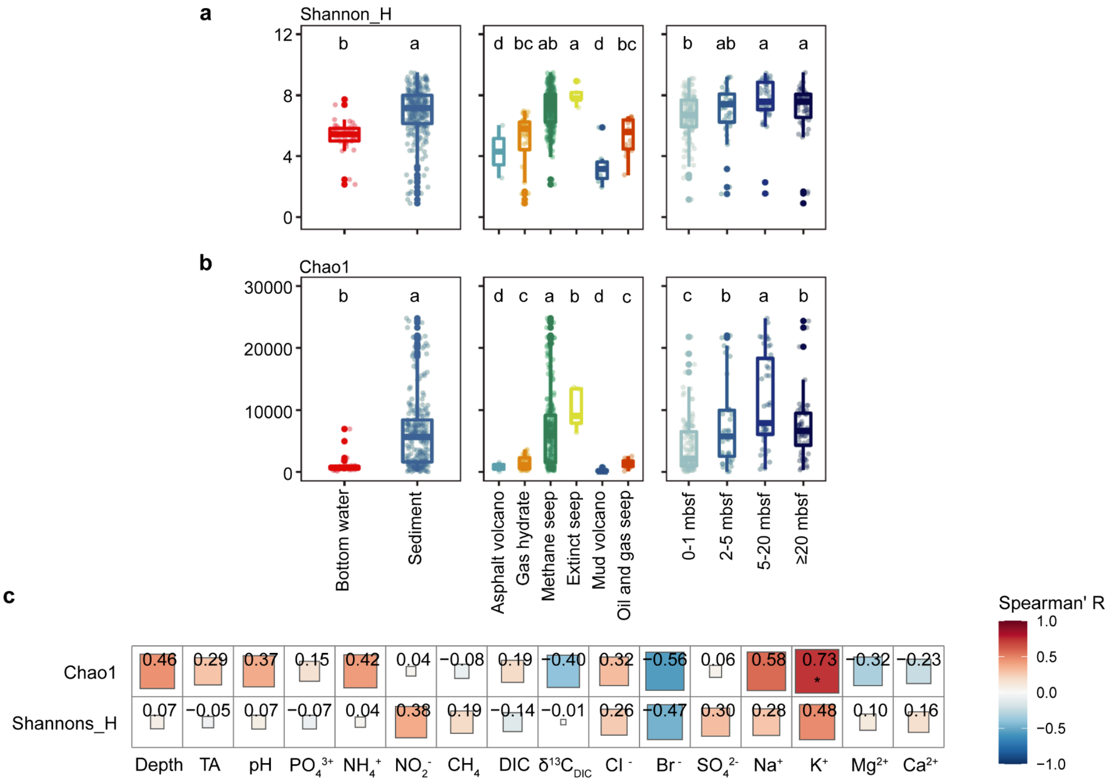


**Figure S4. Viral alpha diversity patterns and environmental correlations in cold seep ecosystems. (a-b)** Viral alpha diversity across cold seep sample types, seep types, and sediment depths. Shannon **(a)** and Chao1 **(b)** indices were used to quantify viral alpha diversity. Letters above boxplots denote statistical groupings based on pairwise Wilcoxon rank-sum tests with Benjamini-Hochberg correction ($P < 0.05$). Groups labeled with different letters show statistically significant differences. mbsf, meters below seafloor. Detailed statistical summaries are provided in **Table S3**. **(c)** Spearman correlation coefficients between viral alpha diversity and environmental parameters, calculated using samples from W07 P1-P20. (*, $P < 0.05$; **, $P < 0.01$). Detailed statistical summaries are provided in **Table S17**.

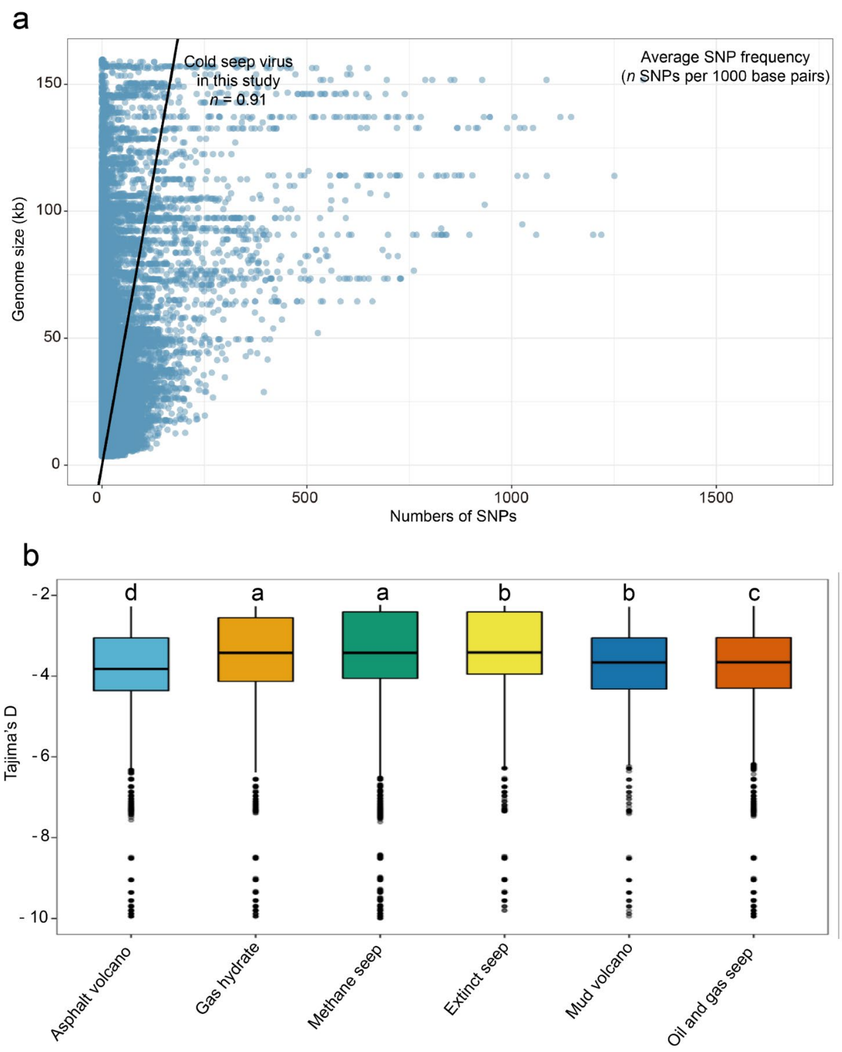


**Figure S5. Microdiversity patterns in the Global Cold Seep Virome (GCSV). (a)** Relationship between genome size and SNP counts in cold seep viral populations. Viral genomes with lengths below the 1st percentile or above the 99th percentile were excluded, and 90,000 genomes were randomly subsampled from the remaining dataset for visualization. Each point represents an individual viral population, and the solid line indicates the expected relationship based on the average SNP density (SNPs/kb). **(b)** Distribution of Tajima's D values for viral genes across six seep types. Letters above boxplots denote statistical groupings based on pairwise Wilcoxon rank-sum tests with Benjamini-Hochberg correction ($P < 0.05$). Groups labeled with different letters show statistically significant differences. Detailed statistics are provided in **Table S3**.

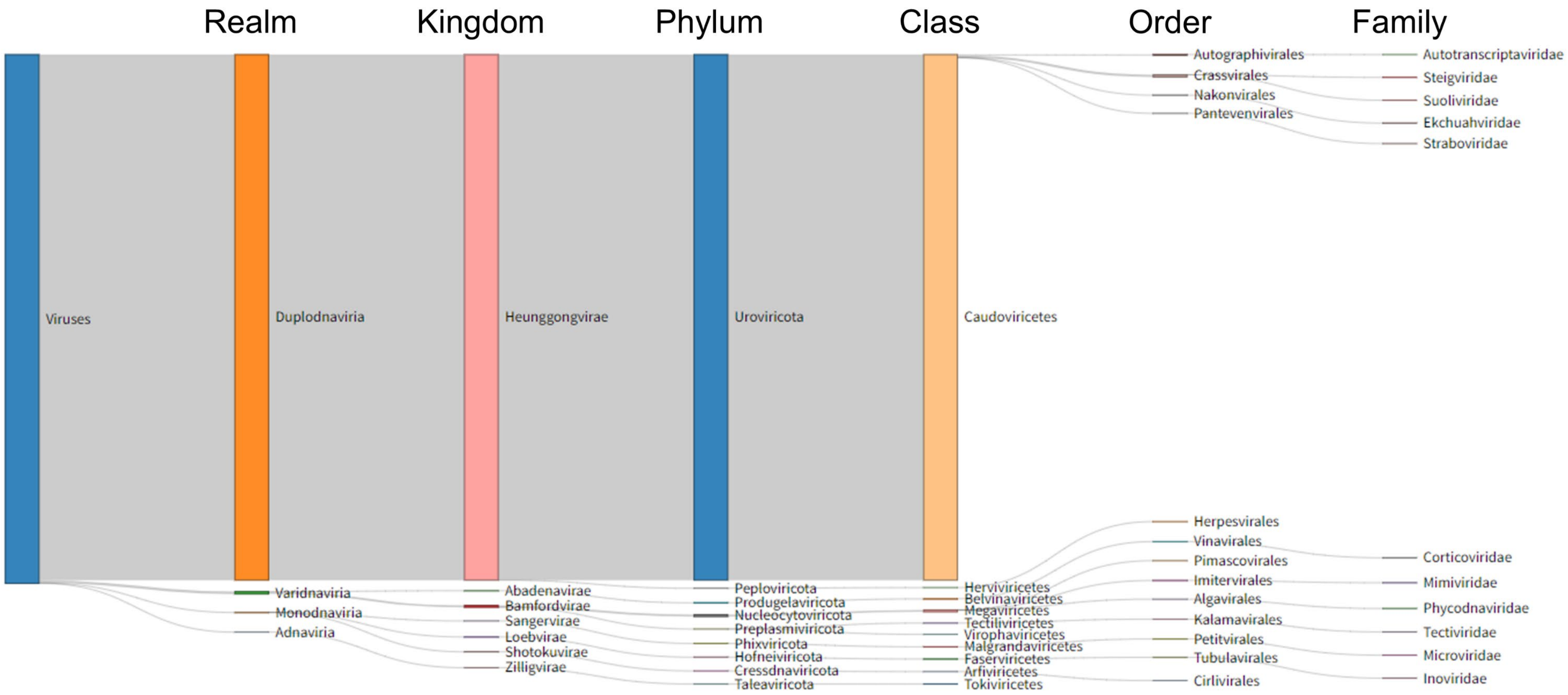


**Figure S6. Hierarchical taxonomic classification of vOTUs in the GCSV.** Sankey diagram showing the taxonomic assignment of vOTUs from the realm to the family level. Line widths represent the relative number of vOTUs assigned at each taxonomic rank. Detailed classification statistics are provided in **Table S4**.

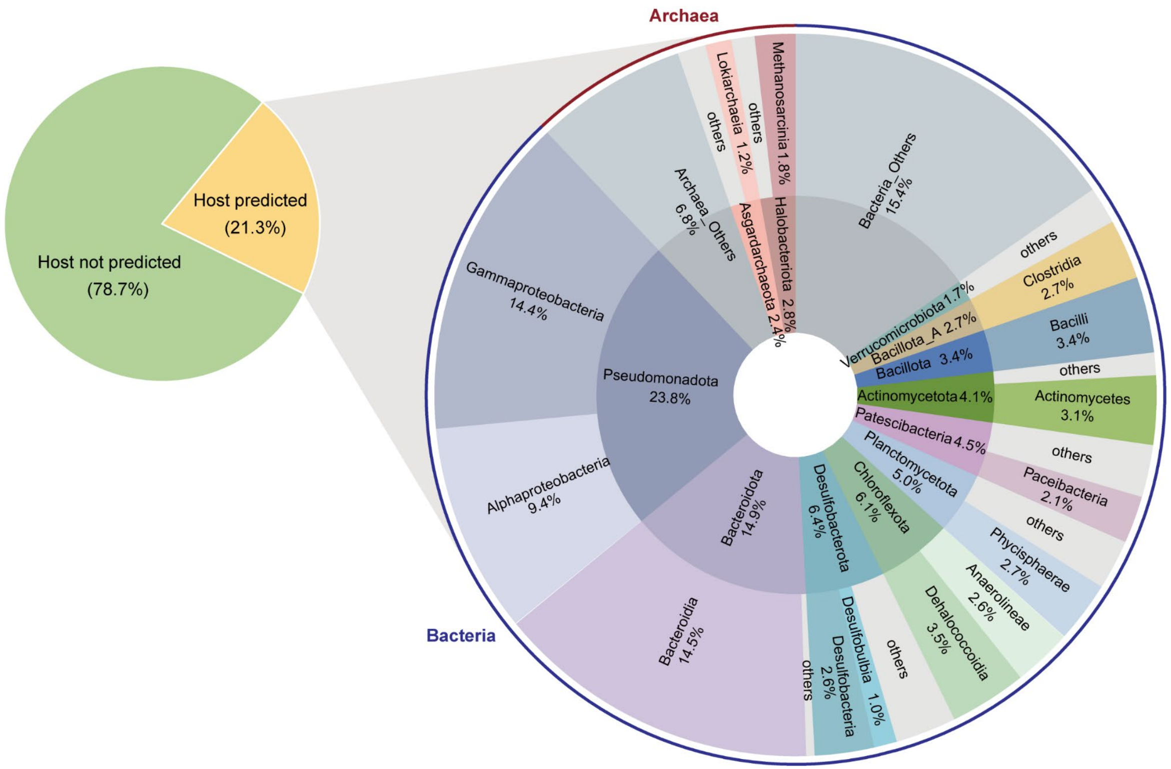


**Figure S7. Predicted viral host taxonomy of vOTUs.** The pie chart (left) shows the proportion of vOTUs with assigned microbial hosts (yellow) and without host assignments (green). The circular plot (right) illustrates the taxonomic distribution of predicted hosts, organized by domain (outer ring), phylum (inner ring), and class (middle ring). Percentages in the circular plot represent the quantity of host genomes associated with each taxonomic group. Bacterial hosts are shown in blue and archaeal hosts in red. Detailed data are provided in **Table S5**.

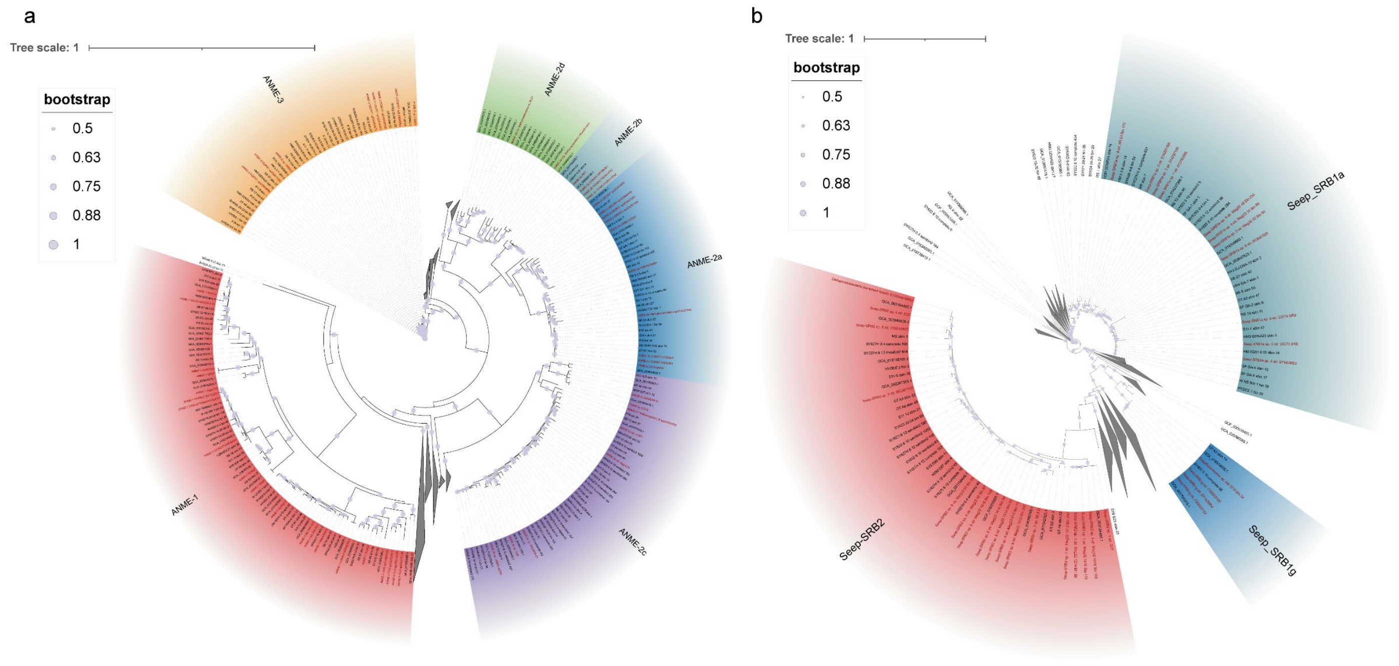


**Figure S8. Phylogenomic reconstruction of anaerobic methanotrophic archaea (ANME) and syntrophic sulfate-reducing bacteria (SRB). (a)** Maximum-likelihood phylogenomic tree of ANME and related *Halobacteriota* MAGs recovered in this study (black) and reference ANME genomes (red), based on a concatenated alignment of 53 conserved archaeal single-copy marker genes. **(b)** Maximum-likelihood phylogenomic tree of syntrophic SRB MAGs affiliated with *Desulfobacterota* from this study (black) and reference SRB genomes (red), reconstructed from 120 conserved bacterial single-copy marker genes. For both trees, node size indicates bootstrap support, and scale bars represent the number of substitutions per site.

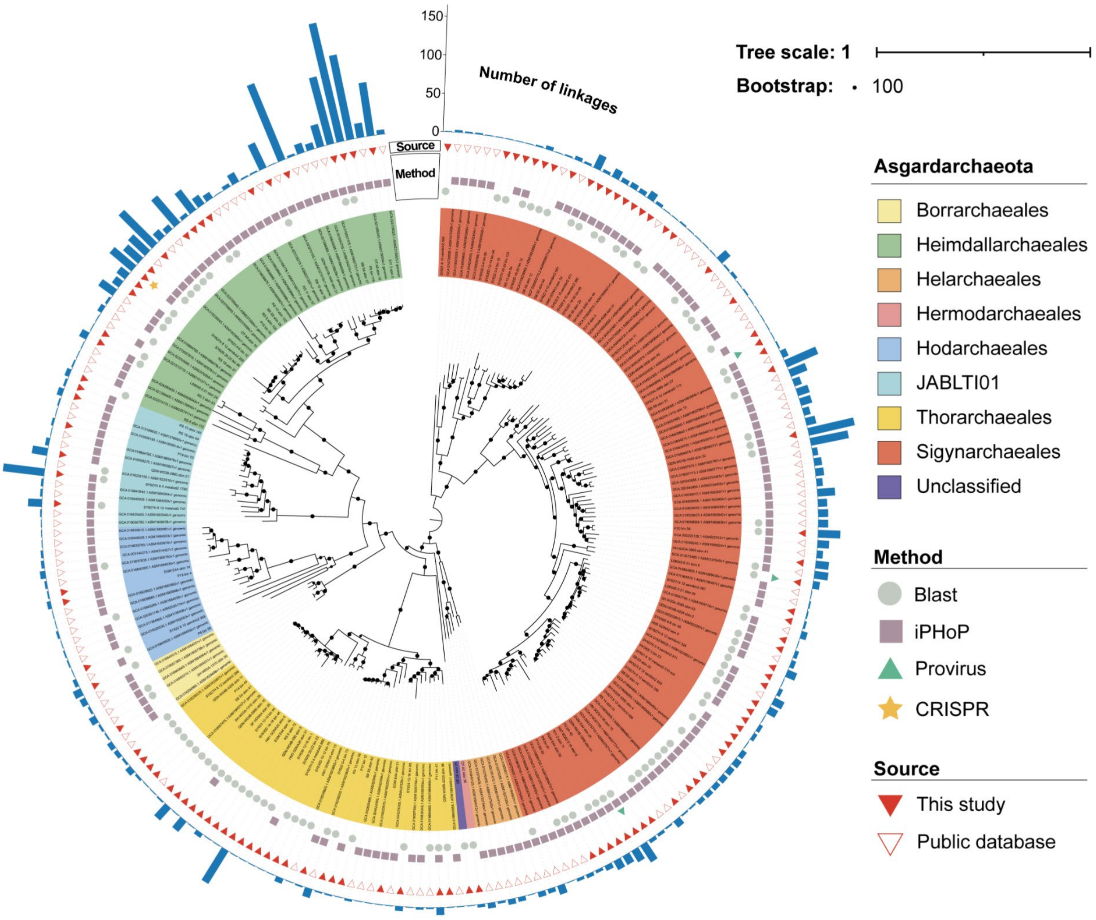


**Figure S9. Diversity of *Asgardarchaeota* genomes and associated viral linkages.** Maximum-likelihood phylogenomic tree of *Asgardarchaeota* genomes inferred from a concatenated alignment of 53 conserved archaeal single-copy marker genes. Background shading indicates the taxonomic order. From inside to outside, the tree rings show: (i) method of host-linkage inference; (ii) genome source; and (iii) number of vOTUs linked to each host genome. The scale bar indicates the number of substitutions per site and the node size is shown only for nodes with bootstrap support of 100.

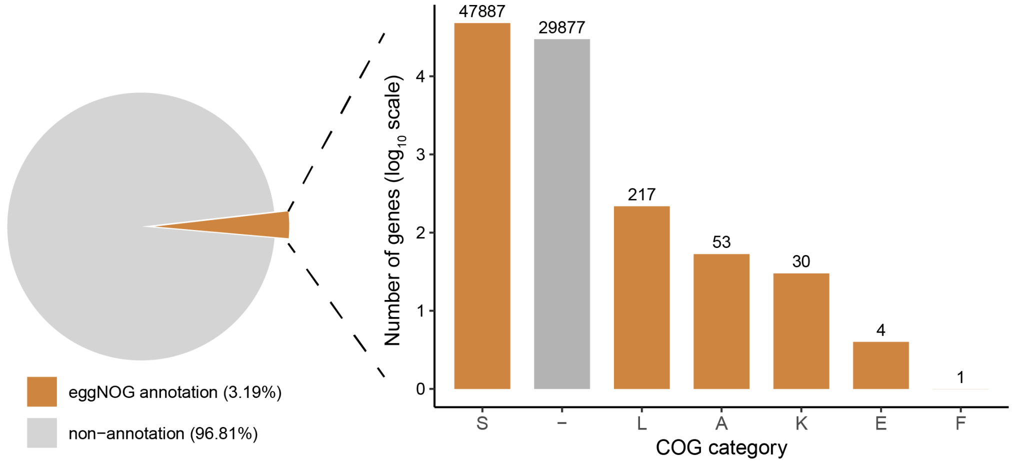


**Figure S10. Protein annotations generated using eggNOG-mapper.** The left panel shows the proportion of viral proteins that received functional annotations from eggNOG-mapper. The right panel displays the distribution of annotated proteins across COG functional categories. COG category labels: S, Function unknown; L, Replication, recombination, and repair; A, RNA processing and modification; K, Transcription; E, Amino acid transport and metabolism; F, Nucleotide transport and metabolism; –, No COG assignment.

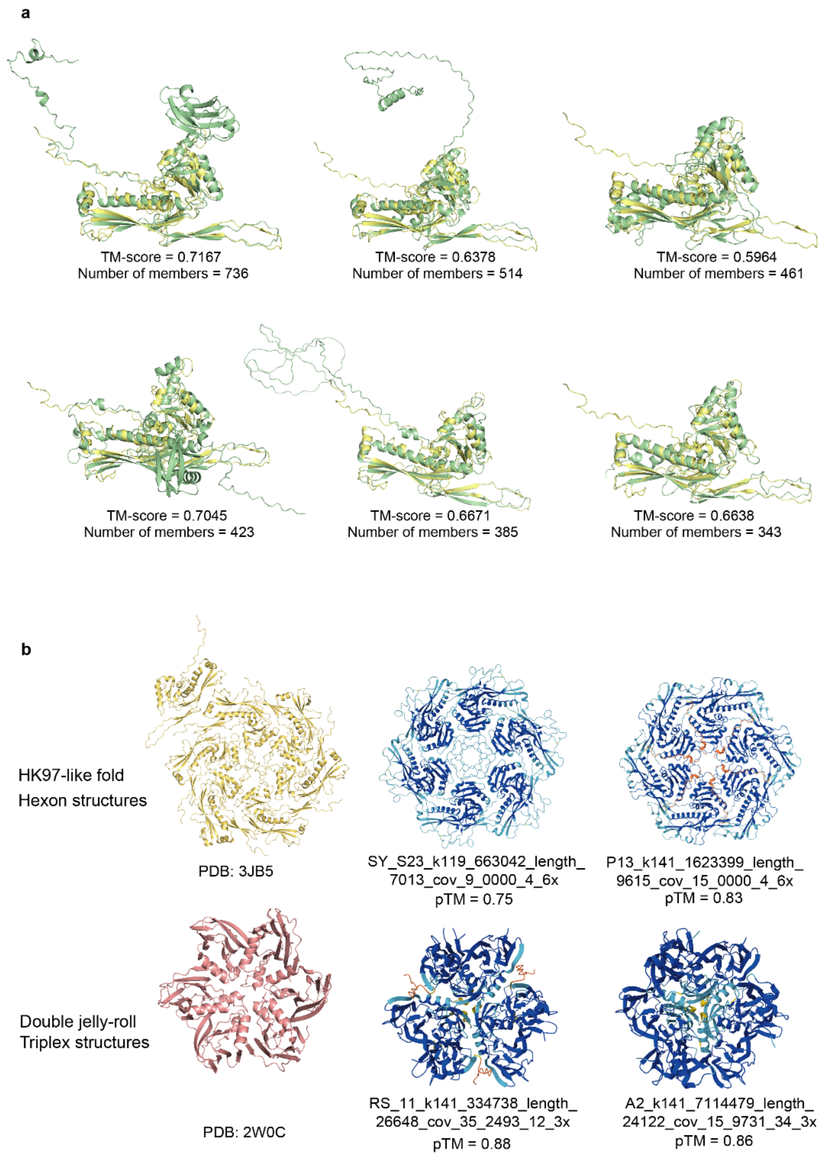


**Figure S11. Structures and multimeric assemblies of major capsid proteins (MCPs). (a)** Representative monomeric MCP structures from the six largest MCP viral protein clusters (vPCs) in the GCSV (green) superimposed onto an HK97-like reference MCP (yellow; PDB: 3JB5, chain A). For each vPC, the TM-score of the structural alignment and the total number of cluster members are shown. **(b)** Predicted multimeric MCP assemblies generated using AlphaFold3. Top: HK97-like hexameric capsomers, including the reference hexon (left; PDB: 3JB5) and two representative GCSV hexon predictions, with corresponding predicted template modeling scores (pTM). Bottom: Double–jelly-roll triplex structures, including the reference triplex protein (left; PDB: 2W0C) and two GCSV triplex predictions, each accompanied by its pTM value.

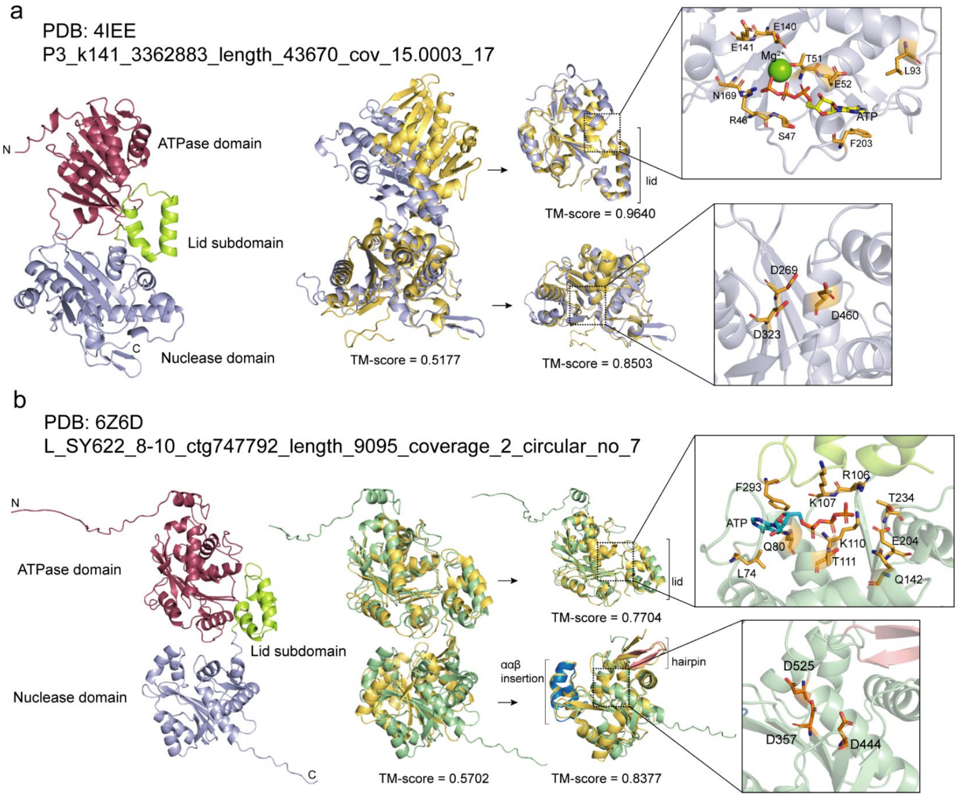


**Figure S12. Domain architecture and active-site conservation of terminase large subunits (TerLs) in the GCSV. (a)** Structural comparison of a GCSV TerL with a reference TerL (PDB: 4IEE). Left: Domain organization of the GCSV TerL, showing the N-terminal ATPase domain, the lid subdomain, and the C-terminal nuclease domain. Middle: Superpositions of the full-length protein and individual domains onto the reference (reference in yellow; GCSV model in green). TM-scores indicate overall and domain-level structural similarity (full-length TM = 0.5177; ATPase TM = 0.9640; nuclease TM = 0.8503). Right: Close-up views of the ATP-binding pocket (with ATP/$Mg^{2+}$ shown) and the RNase H–like nuclease active site. Conserved catalytic and metal-coordinating residues are highlighted. **(b)** Structural comparison of another GCSV TerL with a reference TerL (PDB: 6Z6D). Left: Domain architecture as in panel **(a)**. Middle: Superposition of the full-length model and individual domains (full-length TM = 0.5702; ATPase TM = 0.7704; nuclease TM = 0.8377). The ααβ insertion characteristic of the ATPase domain and the β-hairpin motif in the nuclease domain are indicated. Right: Close-up views of the ATP-binding pocket and nuclease catalytic center with conserved residues shown.

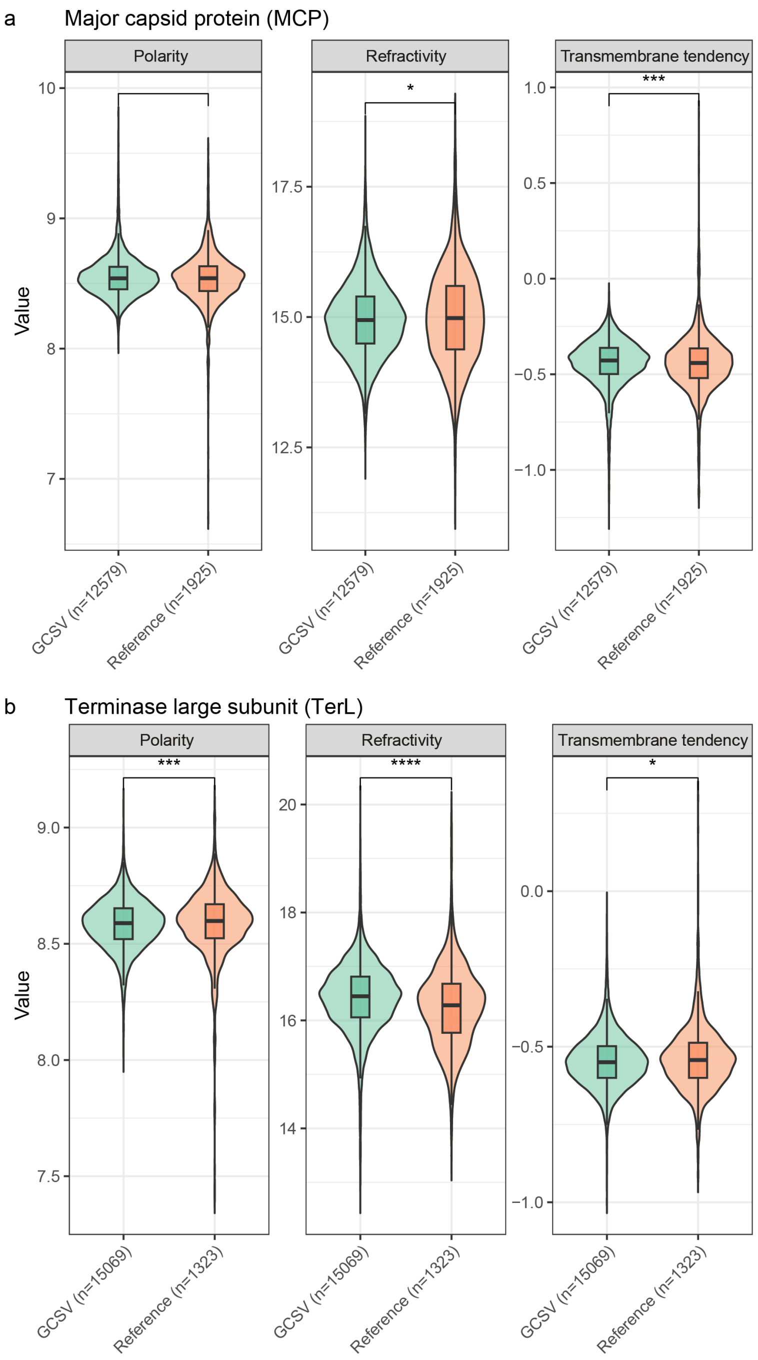


**Figure S13. Physicochemical properties of viral hallmark proteins.** Violin plots summarizing distributions of polarity, refractivity and relative mutability for MCPs **(a)** and TerLs **(b)** from the GCSV (green) compared with reference datasets (orange). Boxplots inside each violin depict the median and interquartile range. *n* indicates the number of proteins in each group. Statistical significance of group differences was evaluated using the Wilcoxon rank-sum test. Significance thresholds are indicated as follows: *$P < 0.05$, **$P < 0.01$, ***$P < 0.001$, ****$P < 0.0001$.

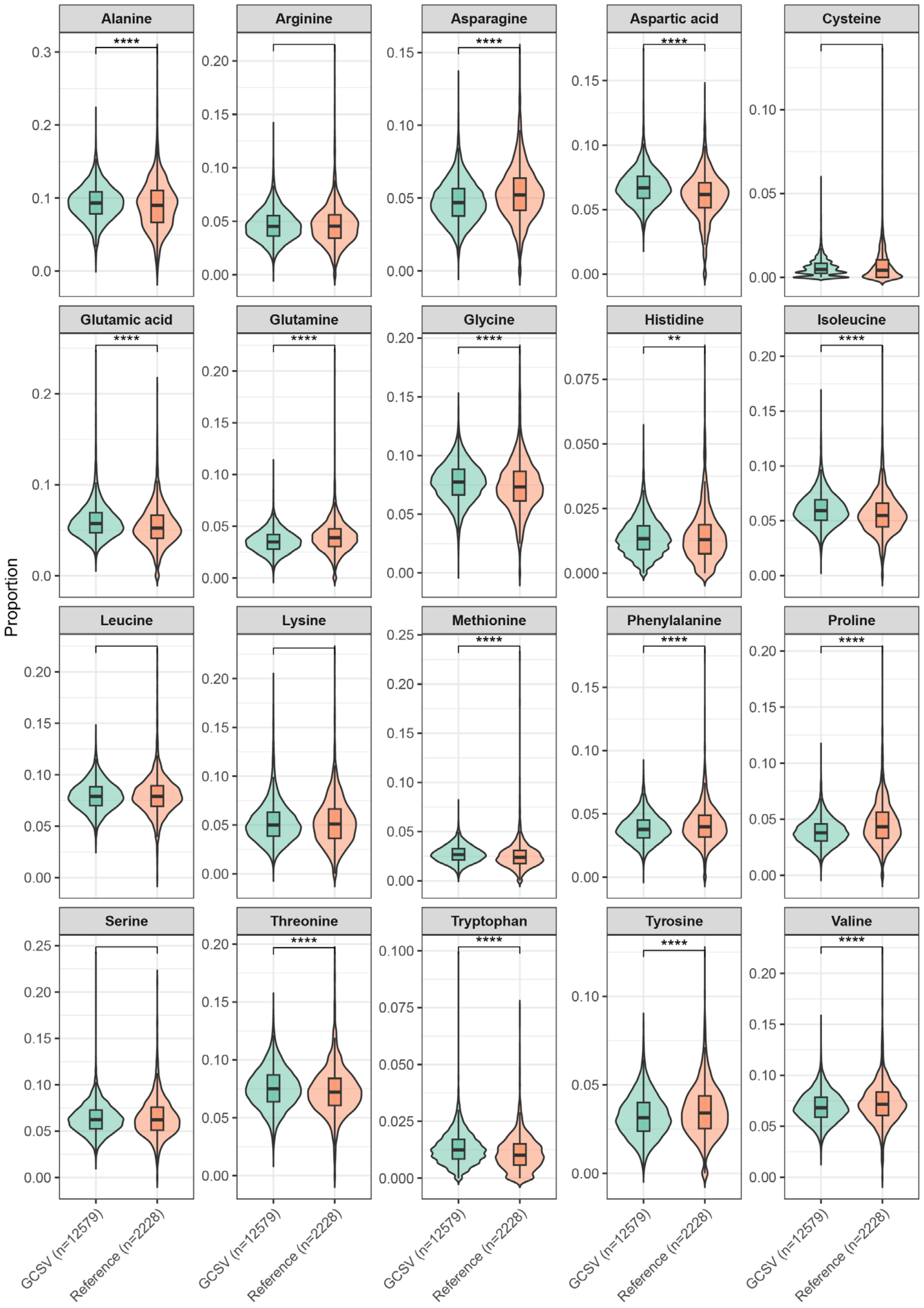


**Figure S14. Amino acid composition of major capsid proteins (MCPs).** Violin plots showing the relative abundance of each amino acid in MCPs from the GCSV (green) compared with reference datasets (orange). Boxplots inside violins represent the median and interquartile range. *n* indicates the number of proteins in each group. Statistical differences between groups were evaluated using the Wilcoxon rank-sum test. Significance thresholds are indicated as follows: *$P < 0.05$, **$P < 0.01$, ***$P < 0.001$, ****$P < 0.0001$. Detailed statistics are provided in **Table S8**.

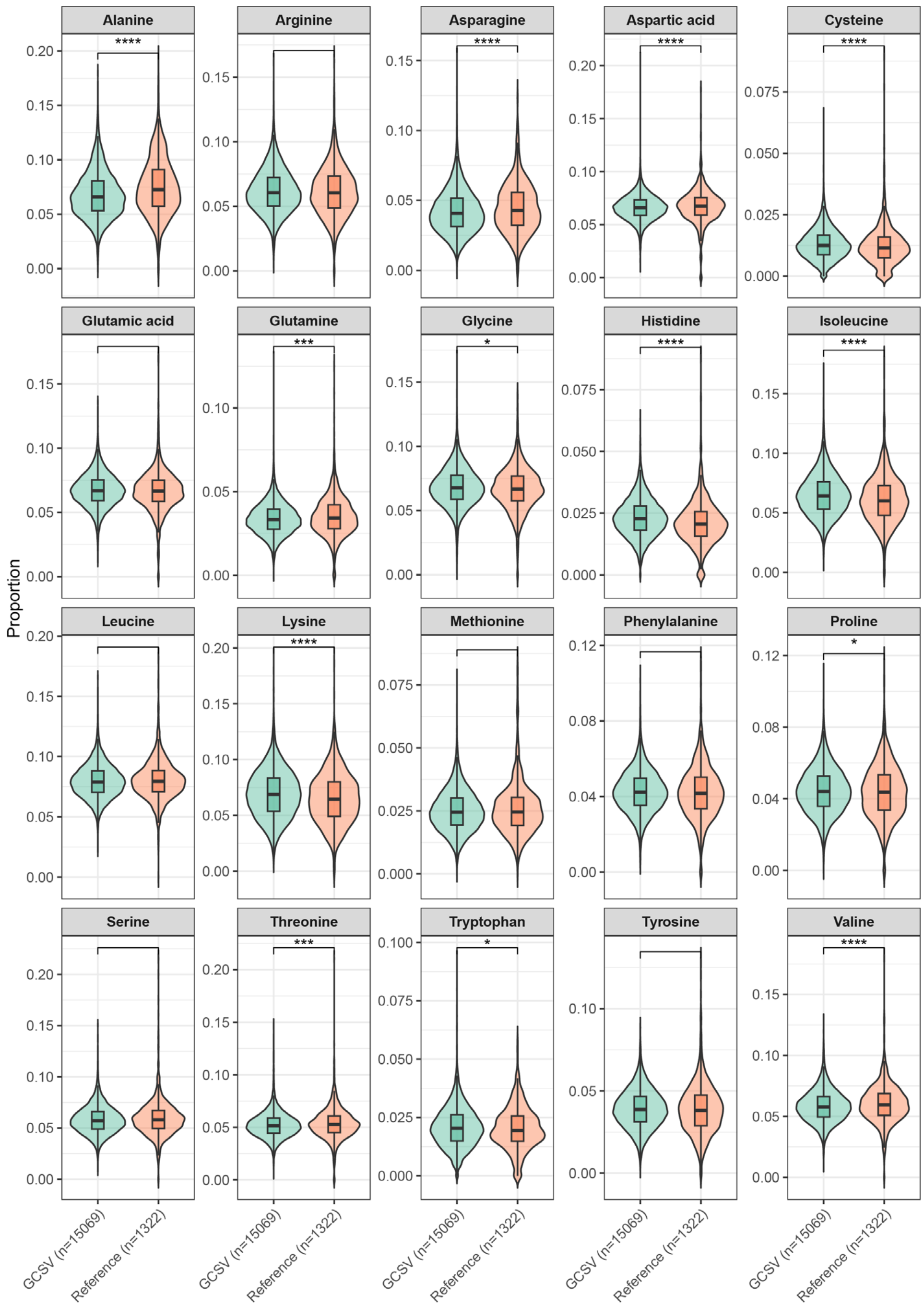


**Figure S15. Amino acid composition of terminase large subunits (TerLs).** Violin plots showing the relative abundance of each amino acid in TerLs from the GCSV (green) compared with reference datasets (orange). Boxplots embedded within violins represent median values and interquartile ranges. *n* indicates the number of proteins in each group. Statistical differences between groups were evaluated using the Wilcoxon rank-sum test. Significance thresholds are indicated as follows: *$P < 0.05$, **$P < 0.01$, ***$P < 0.001$, ****$P < 0.0001$. Detailed statistics are provided in **Table S8**.

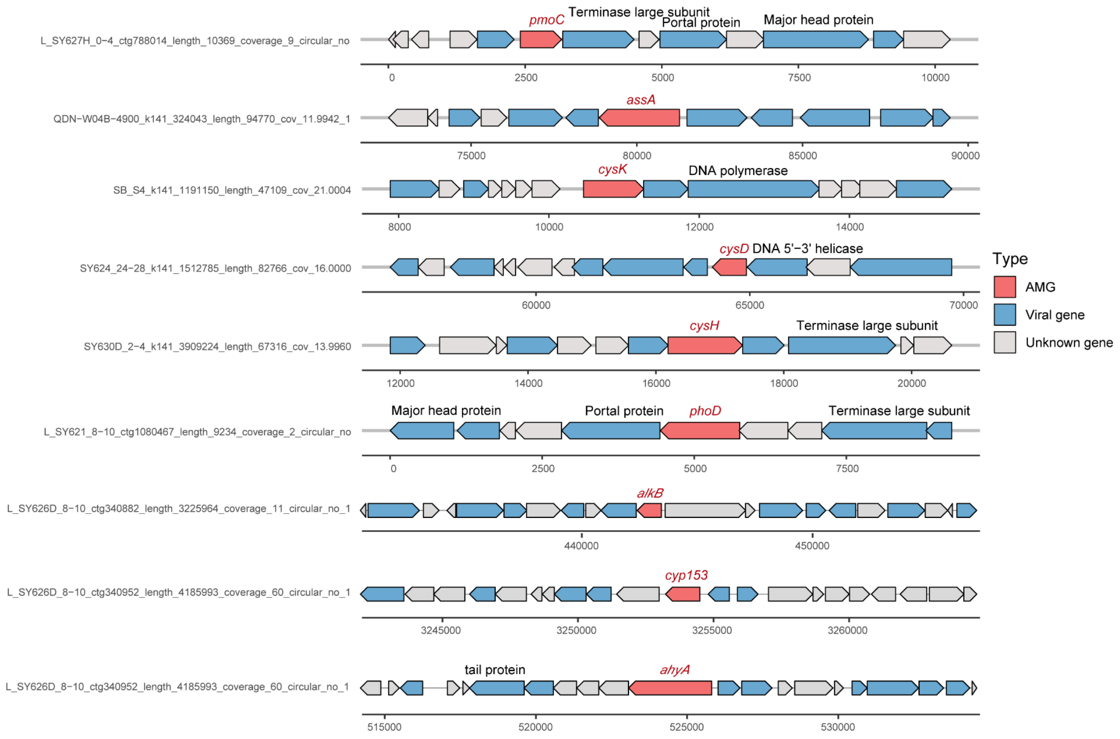


**Figure S16. Genomic organization of viruses encoding key auxiliary metabolic genes (AMGs).** Arrows indicate the position and orientation of predicted open reading frames along each viral genome, with colors denoting gene functional categories derived from Phold annotations, as shown in the legend. AMGs encoded within viral genomes are highlighted in red. Viral hallmark genes are shown in blue, and genes with unknown functions are shown in grey.

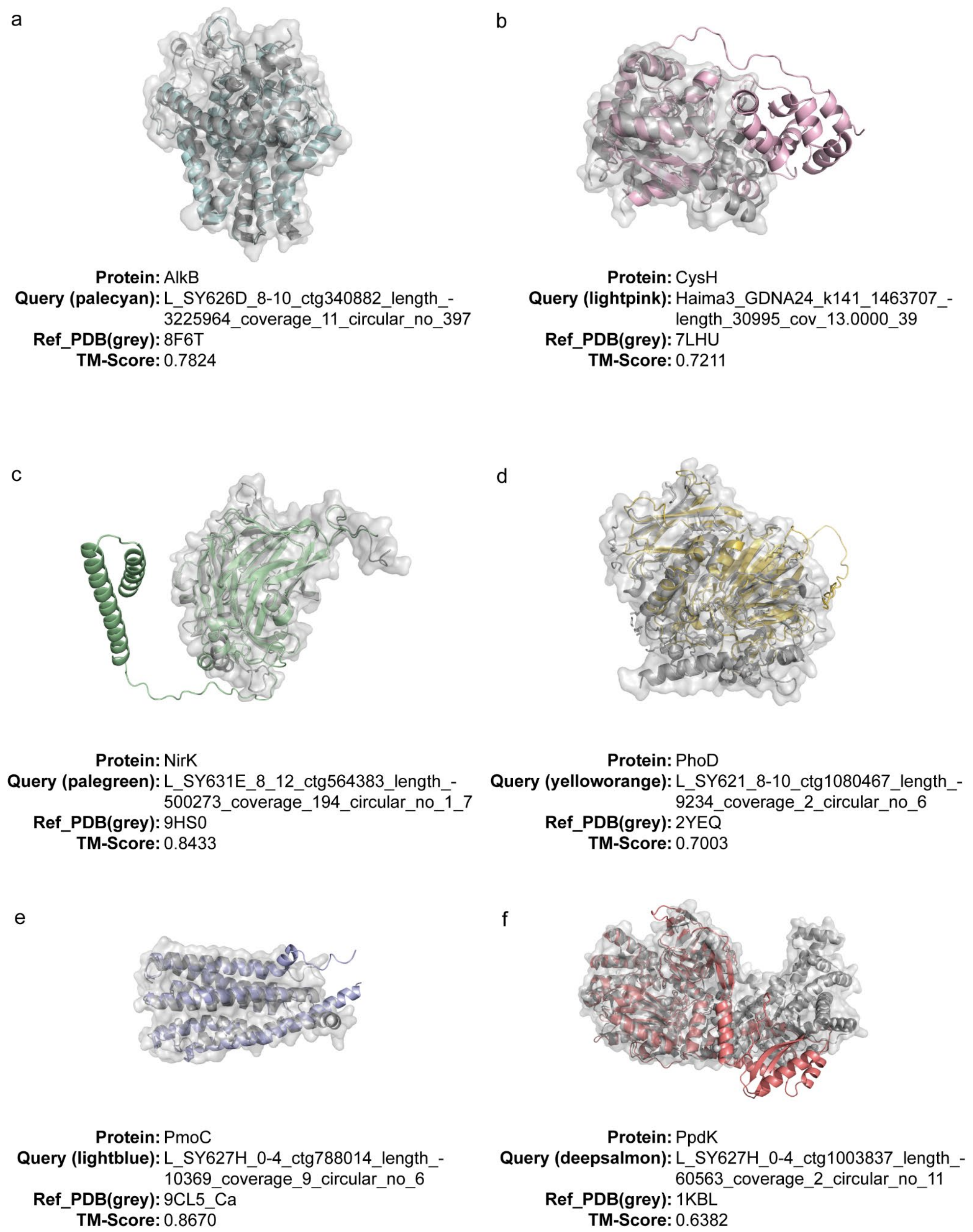


**Figure S17. Structural alignment of representative AMGs from the GCSV with reference proteins.** Structural comparisons are shown for the six AMGs highlighted in **Fig. 4**, including AlkB **(a)**, CysH **(b)**, NirK **(c)**, PhoD **(d)**, PmoC **(e)**, and PpdK **(f).** In each panel, the AMG predicted from the GCSV is displayed in color, while the corresponding reference structure from the PDB is shown in grey. Structural similarity was assessed using TM-align, and the PDB accession number and TM-score for each AMG-reference pair are indicated below the models.

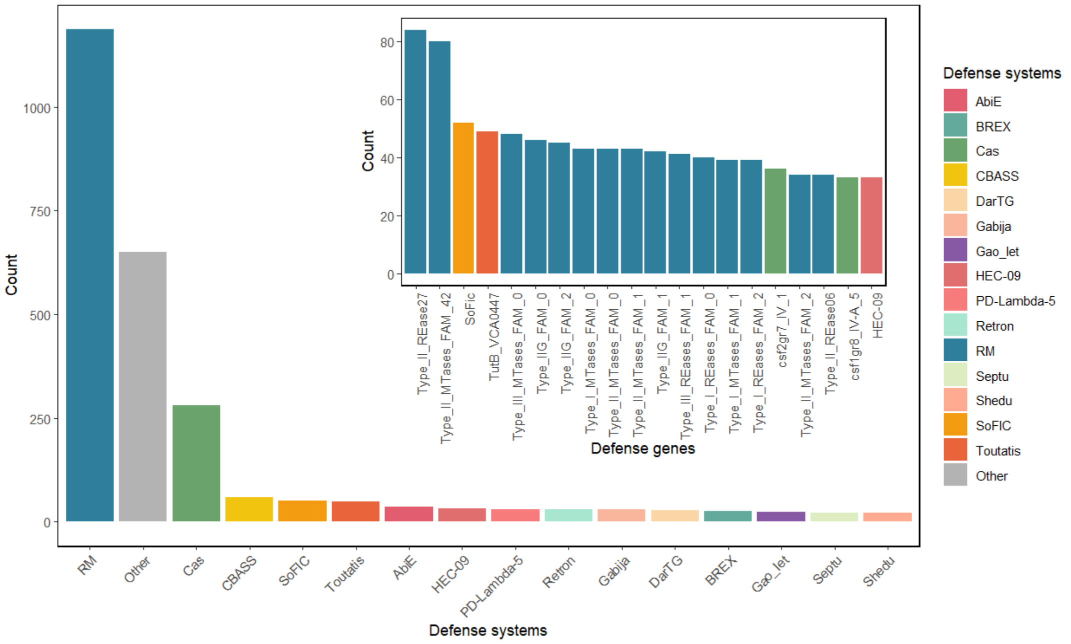


**Figure S18. Distribution of defense genes and systems in the GCSV.** The bar plot (lower left) summarizes the abundance of major prokaryotic defense systems encoded across vPCs in the GCSV, with the 15 most abundant systems individually colored and remaining systems grouped as “Others” (grey). The inset (upper right) displays the top 20 most abundant defense genes, ordered by count. Bar heights represent the number of vPCs containing each defense gene or system.

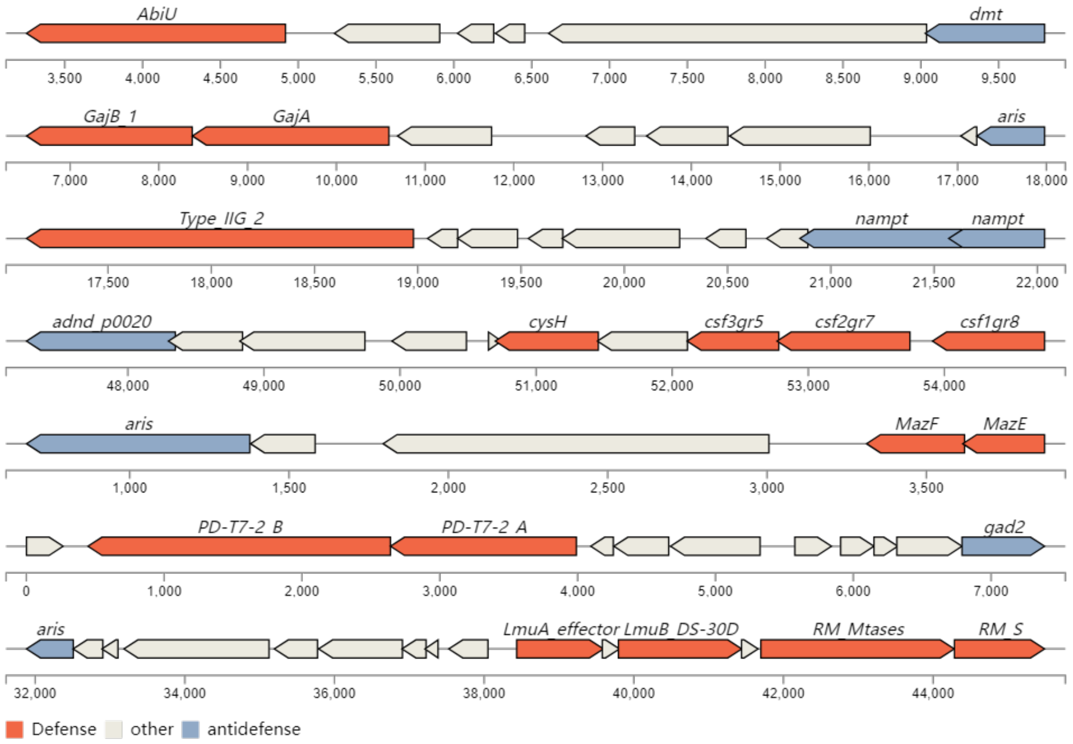


**Figure S19. Co-localization of defense and anti-defense genes within representative viral genomes.** Shown are genomic regions containing both viral defense genes (red) and anti-defense genes (blue), with intervening viral genes depicted in white. Arrows indicate the position and orientation of predicted open reading frames.

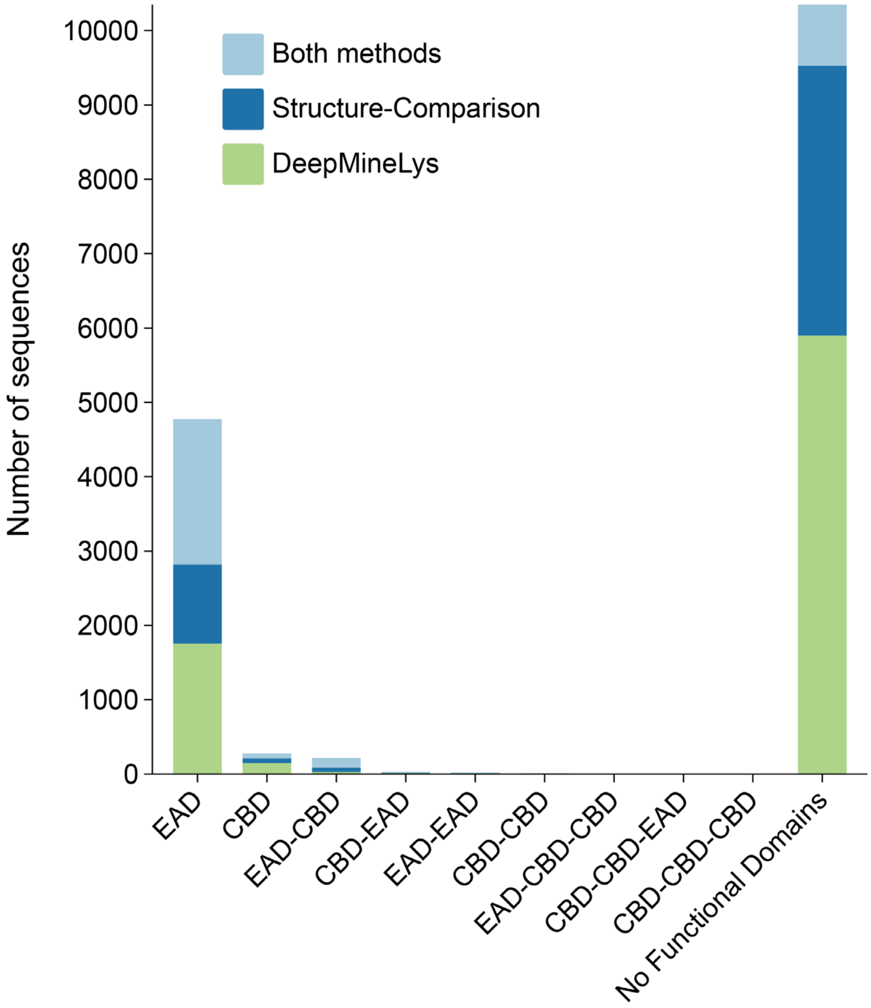


**Figure S20. Classification of predicted lysins based on domain architecture.** Bar plots show the distribution of lysin domain architectures identified in the GCSV. The x-axis lists the detected combinations of enzymatically active domains (EADs) and cell wall binding domains (CBDs), while the y-axis indicates the number of sequences assigned to each category. Colors represent the three lysin identification strategies: DeepMineLys, structure-based comparison, and sequences supported by both methods. The category "No functional domains detected" refers to lysin candidates lacking identifiable EAD or CBD motifs.

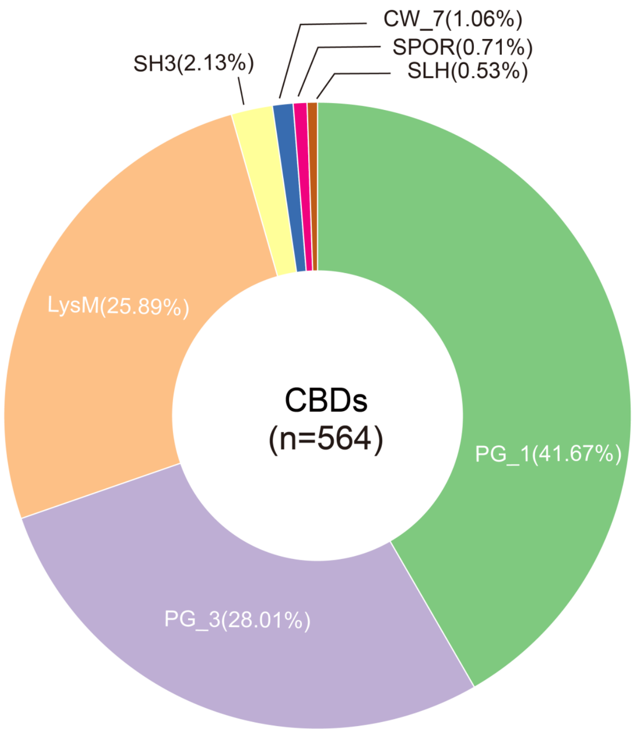


**Figure S21. Classification of C-terminal cell wall-binding domains (CBDs) identified in the GCSV.** The donut chart illustrates the distribution of 564 CBDs across major domain families. PG_1 domains were the most abundant (41.67%), followed by PG_3 (28.01%) and LysM (25.89%). Less represented CBD types included SH3 (2.13%), CW_7 (1.06%), SPOR (0.71%), and SLH (0.53%).

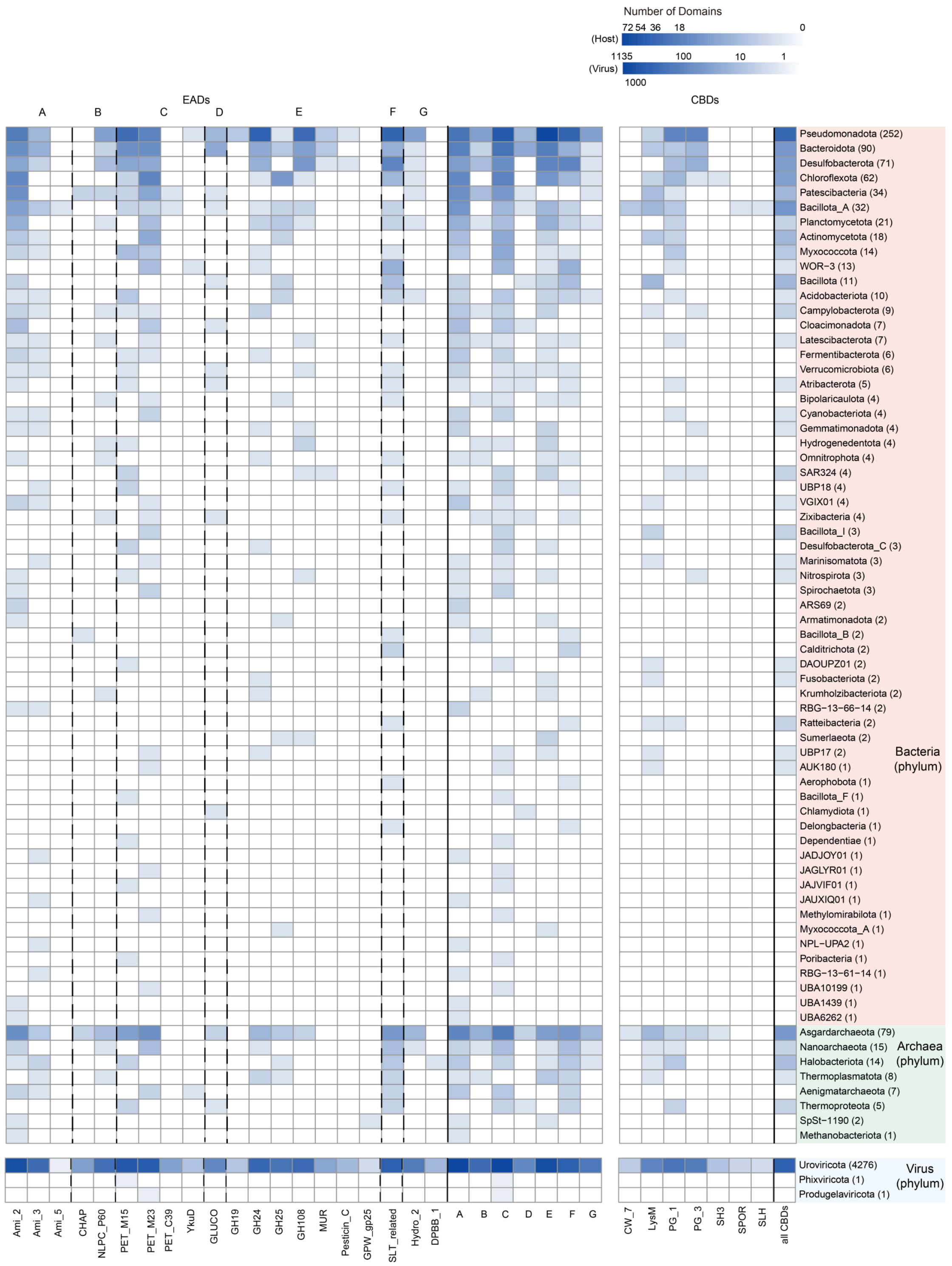


**Figure S22. Distribution of enzymatically active domains (EADs) and cell wall–binding domains (CBDs) across viral host (top) and viral (bottom) phyla.** The heatmap shows the $log_{10}$-transformed abundance of each domain type, with EADs grouped into seven families (A-G) and CBDs summarized on the right. Numbers in parentheses indicate the total number of sequences detected in each phylum.

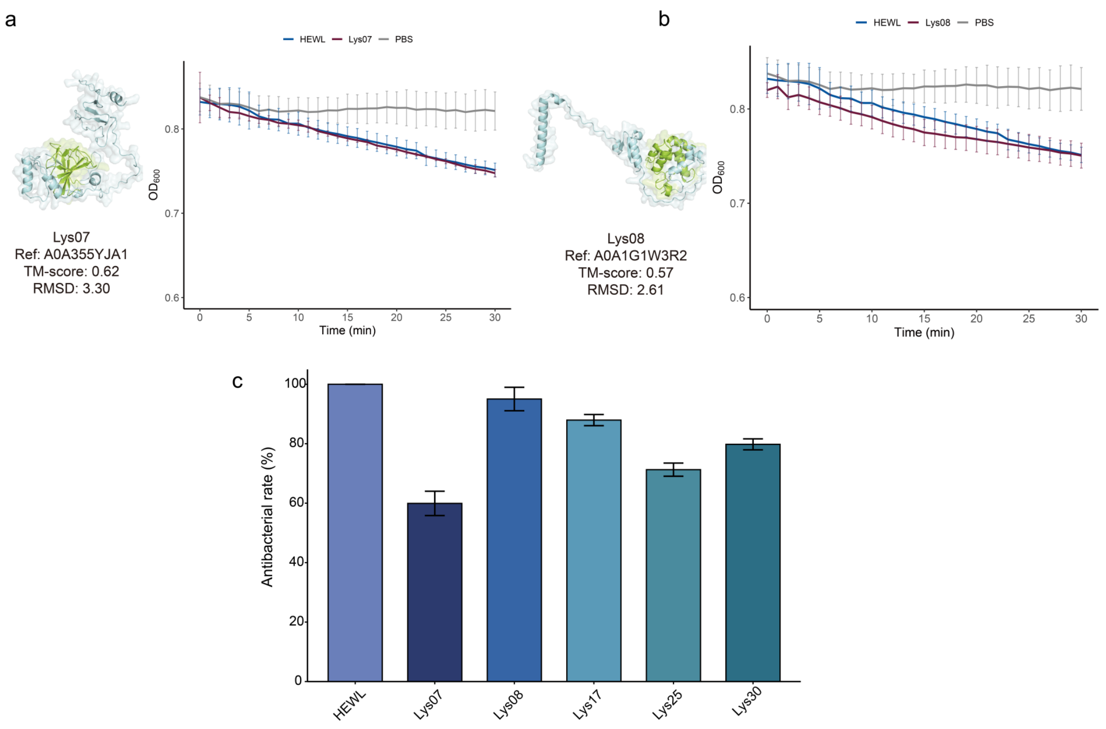


**Figure S23. Structural comparison and functional validation of additional GCSV lysins. (a-b)** Left: Structural alignments of two representative lysins from the GCSV with their closest reference proteins. Reference structures are shown in grey and GCSV lysins in sky blue. For each pair, the reference protein identifier, TM-score, and RMSD are provided. Right: Time-course assays measuring the decrease in $OD_{600}$ of *Bacillus subtilis* ATCC 6051 following treatment with each lysin. HEWL (hen egg white lysozyme) served as a positive control, and PBS as a negative control. Values represent mean ± SD from three independent experiments. **(c)** Antibacterial activities of five GCSV lysins and HEWL against *B. subtilis* ATCC 6051. After incubation with lysins, mixtures were plated on LB agar and colony-forming units were quantified after overnight incubation at 37°C. Error bars denote standard deviations from triplicate assays (n = 3).

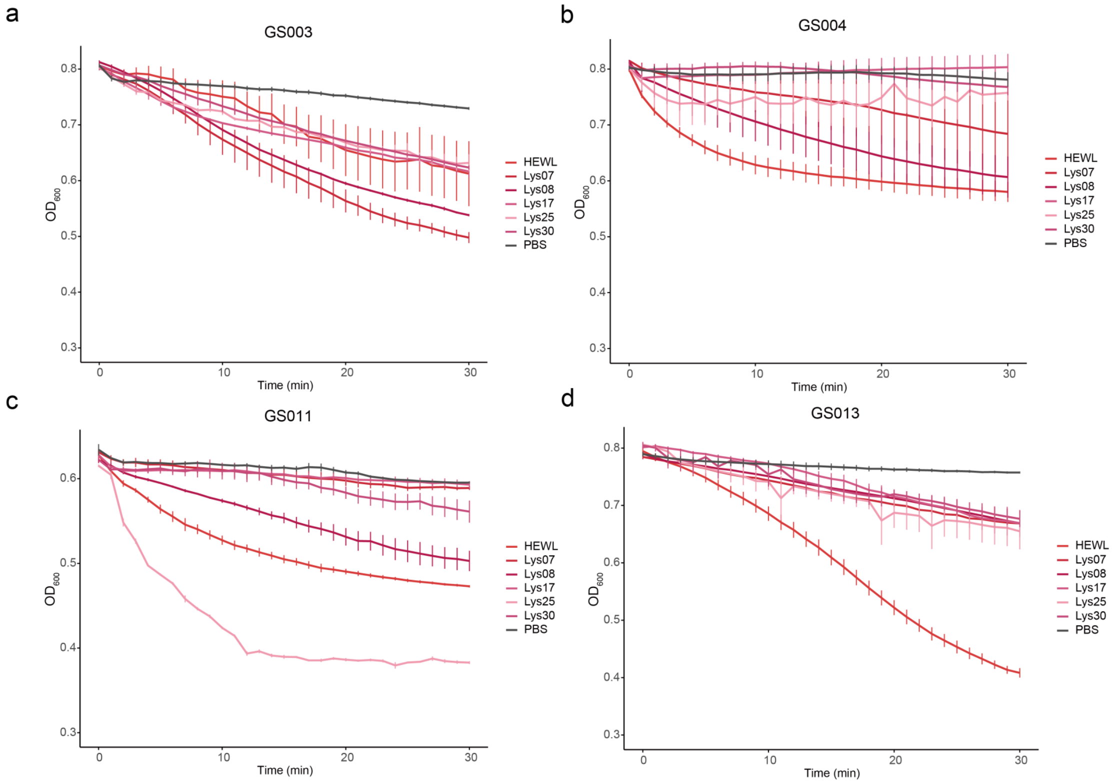


**Figure S24. Lytic activity of GCSV lysins against four deep-sea-derived bacterial strains.** Antibacterial activities of five GCSV lysins and HEWL against *Bacillus tequilensis* GS003 (**a**), *Bacillus altitudinis* GS004 (**b**), *Mesobacillus thioparans* GS011 (**c**), or *Bacillus tequilensis* GS013 (**d**). Against GS003 and GS011, several of these lysins exhibited more potent lytic activity compared to the positive control HEWL. In contrast, their lytic activity was less pronounced than that of HEWL when tested against GS004 and GS013. Values represent mean ± SD from three independent experiments.

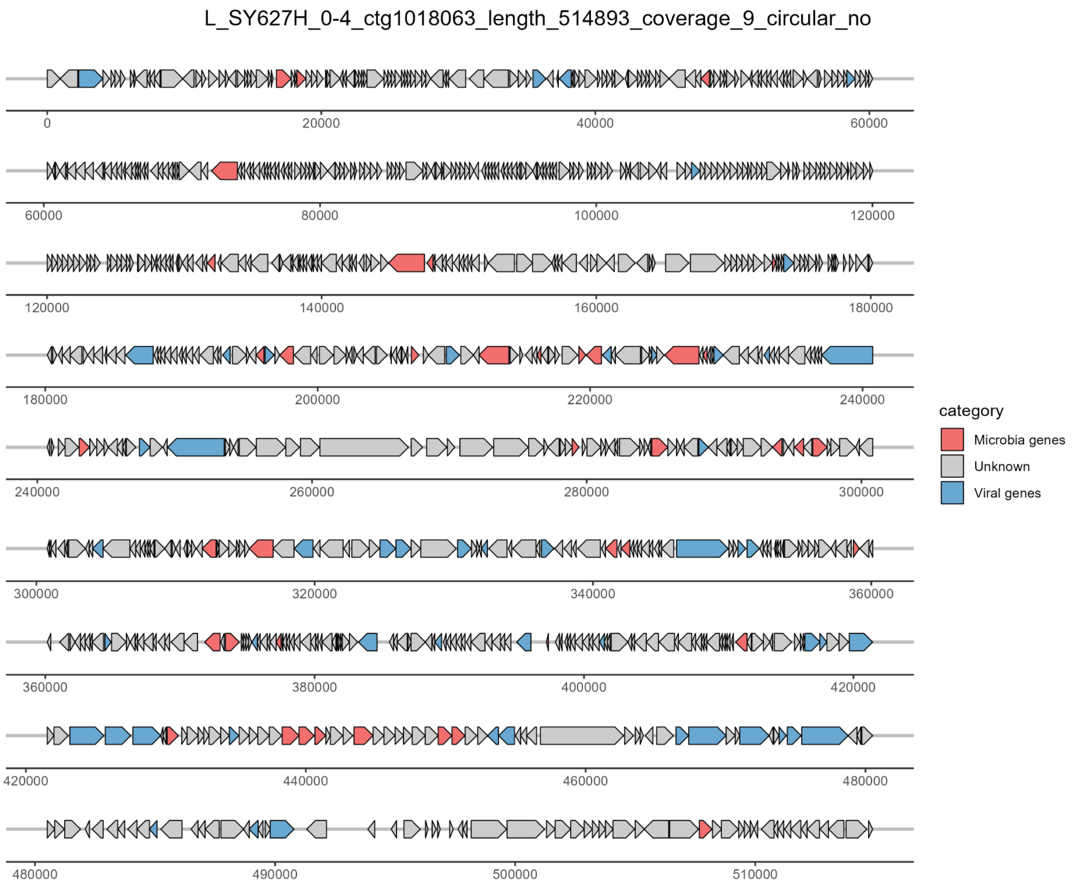
L_SY627H_0-4_ctg1018063_length_514893_coverage_9_circular_no
0
20000
40000
60000
80000
100000
120000
140000
160000
180000
200000
220000
240000
260000
280000
300000
320000
340000
360000
380000
400000
420000
440000
460000
480000
490000
500000
510000
category
Microbia genes
Unknown
Viral genes

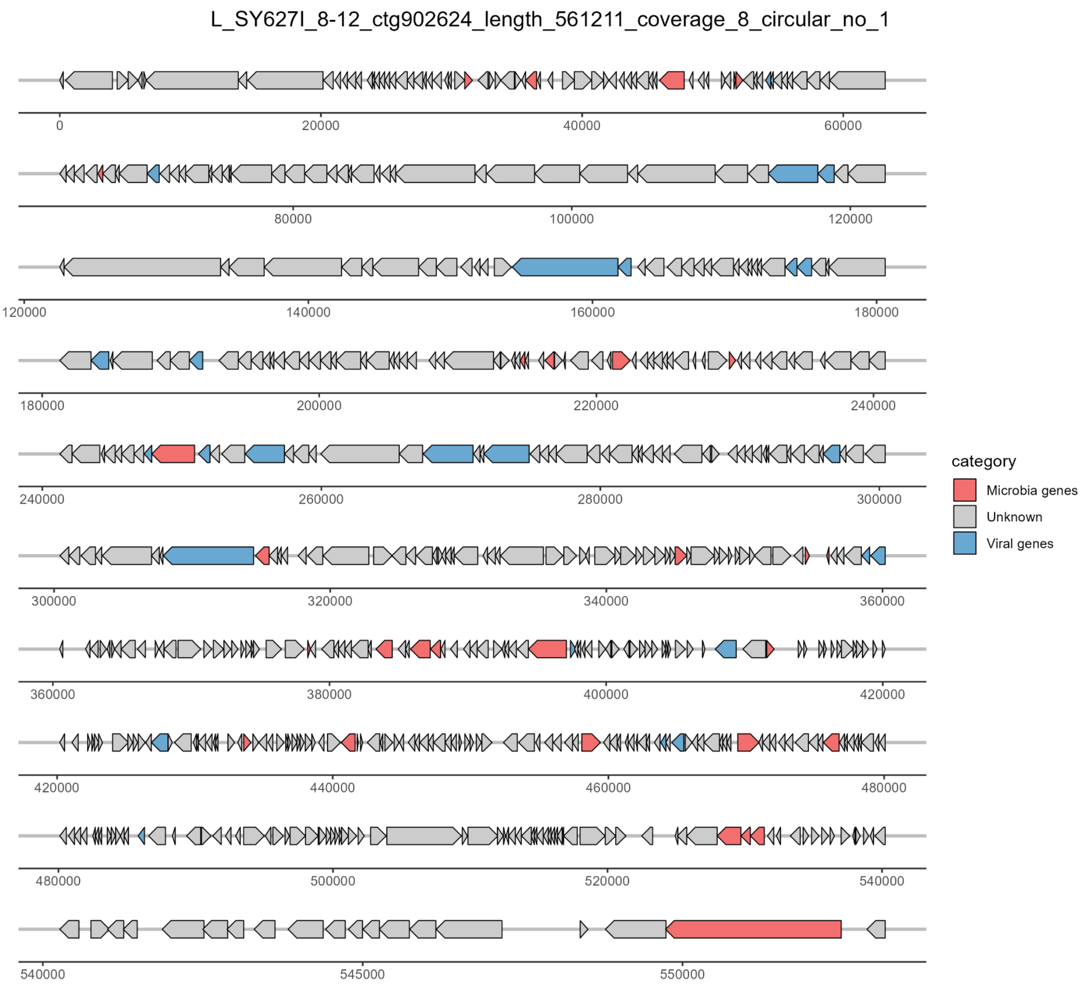
L_SY627I_8-12_ctg902624_length_561211_coverage_8_circular_no_1
0
20000
40000
60000
80000
100000
120000
120000
140000
160000
180000
180000
200000
220000
240000
240000
260000
280000
300000
300000
320000
340000
360000
360000
380000
400000
420000
420000
440000
460000
480000
480000
500000
520000
540000
540000
545000
550000
category
Microbia genes
Unknown
Viral genes

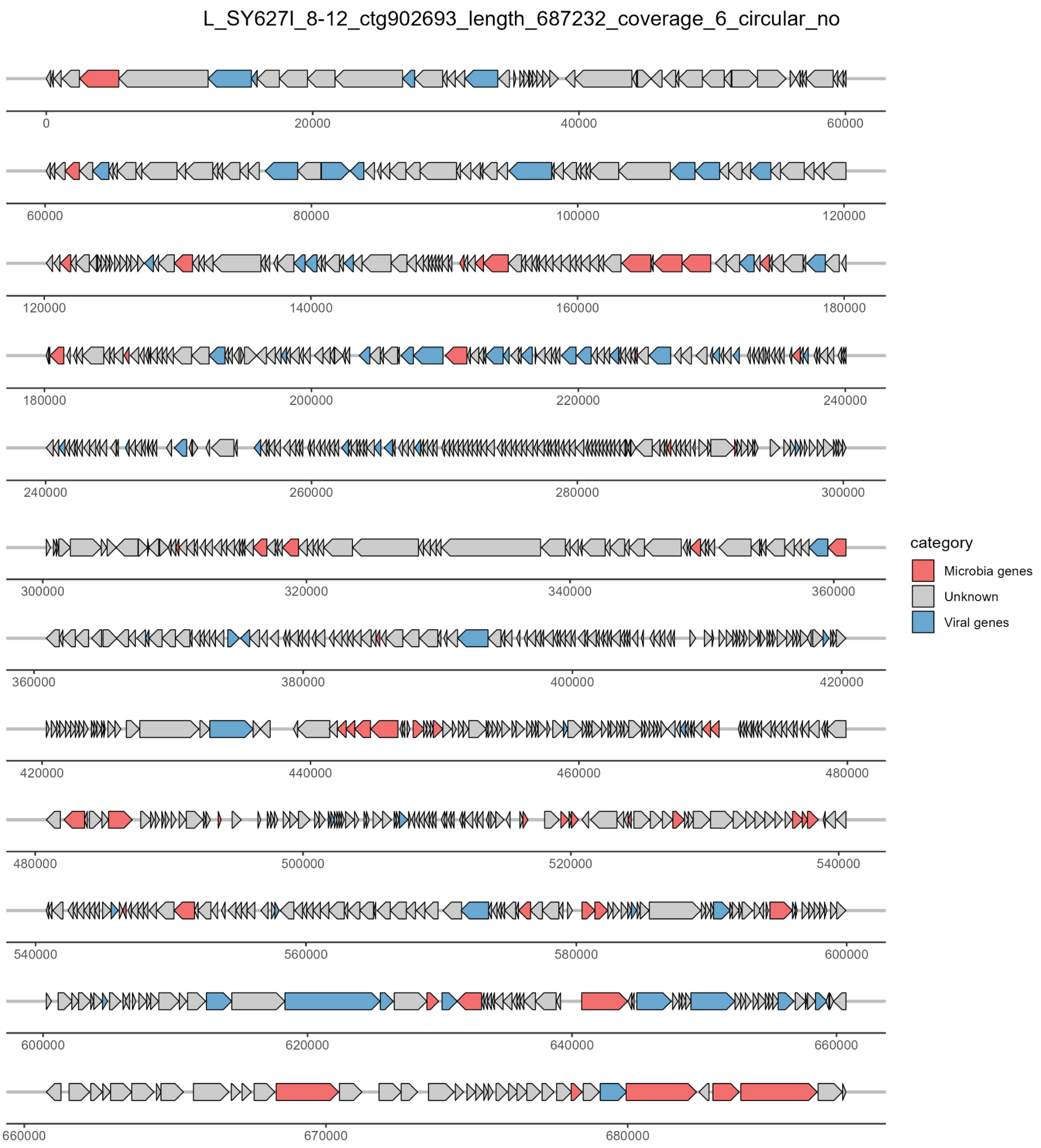

L_SY627l_8-12_ctg902693_length_687232_coverage_6_circular_no
0
20000
40000
60000
80000
100000
120000
140000
160000
180000
200000
220000
240000
260000
280000
300000
320000
340000
360000
380000
400000
420000
440000
460000
480000
500000
520000
540000
560000
580000
600000
620000
640000
660000
670000
680000
category
Microbia genes
Unknown
Viral genes

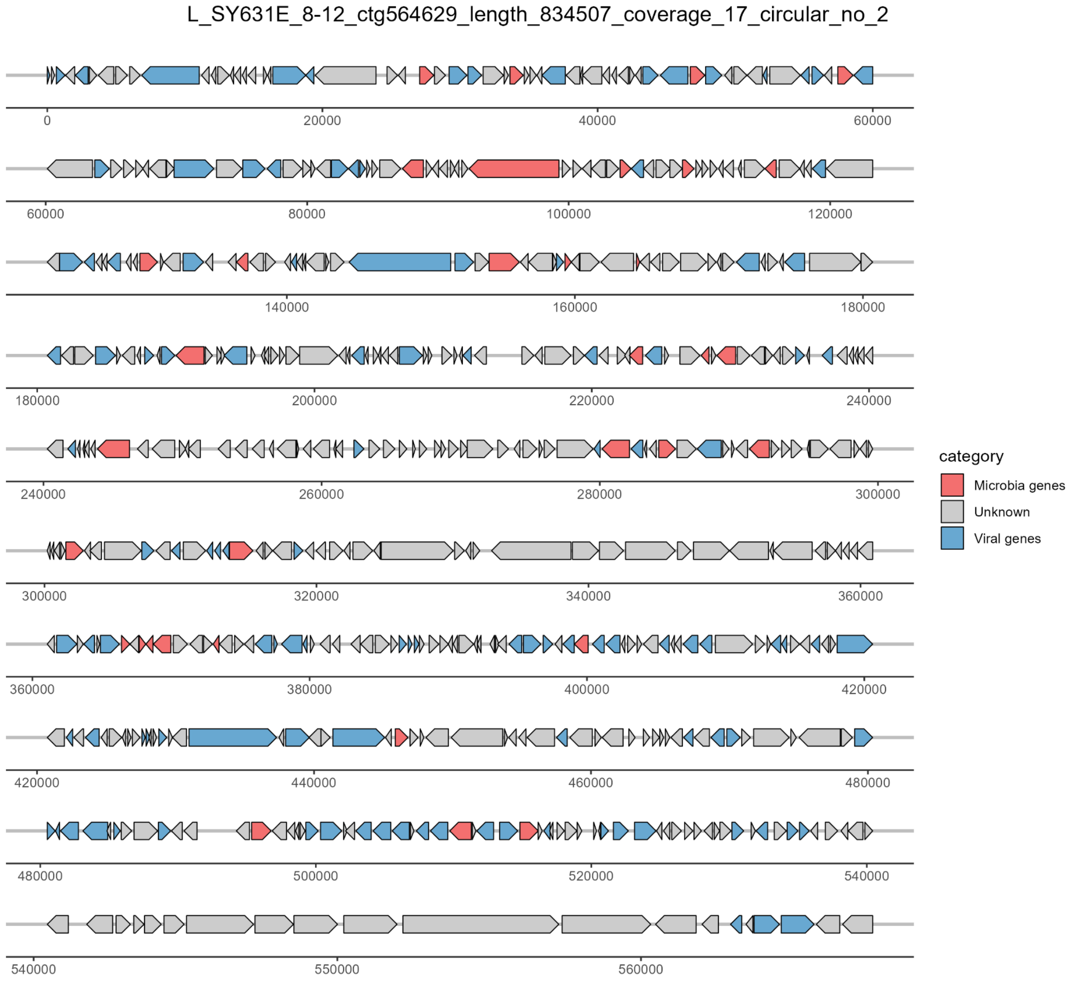
L_SY631E_8-12_ctg564629_length_834507_coverage_17_circular_no_2
0
20000
40000
60000
60000
80000
100000
120000
140000
160000
180000
180000
200000
220000
240000
240000
260000
280000
300000
300000
320000
340000
360000
360000
380000
400000
420000
420000
440000
460000
480000
480000
500000
520000
540000
540000
550000
560000
category
Microbia genes
Unknown
Viral genes

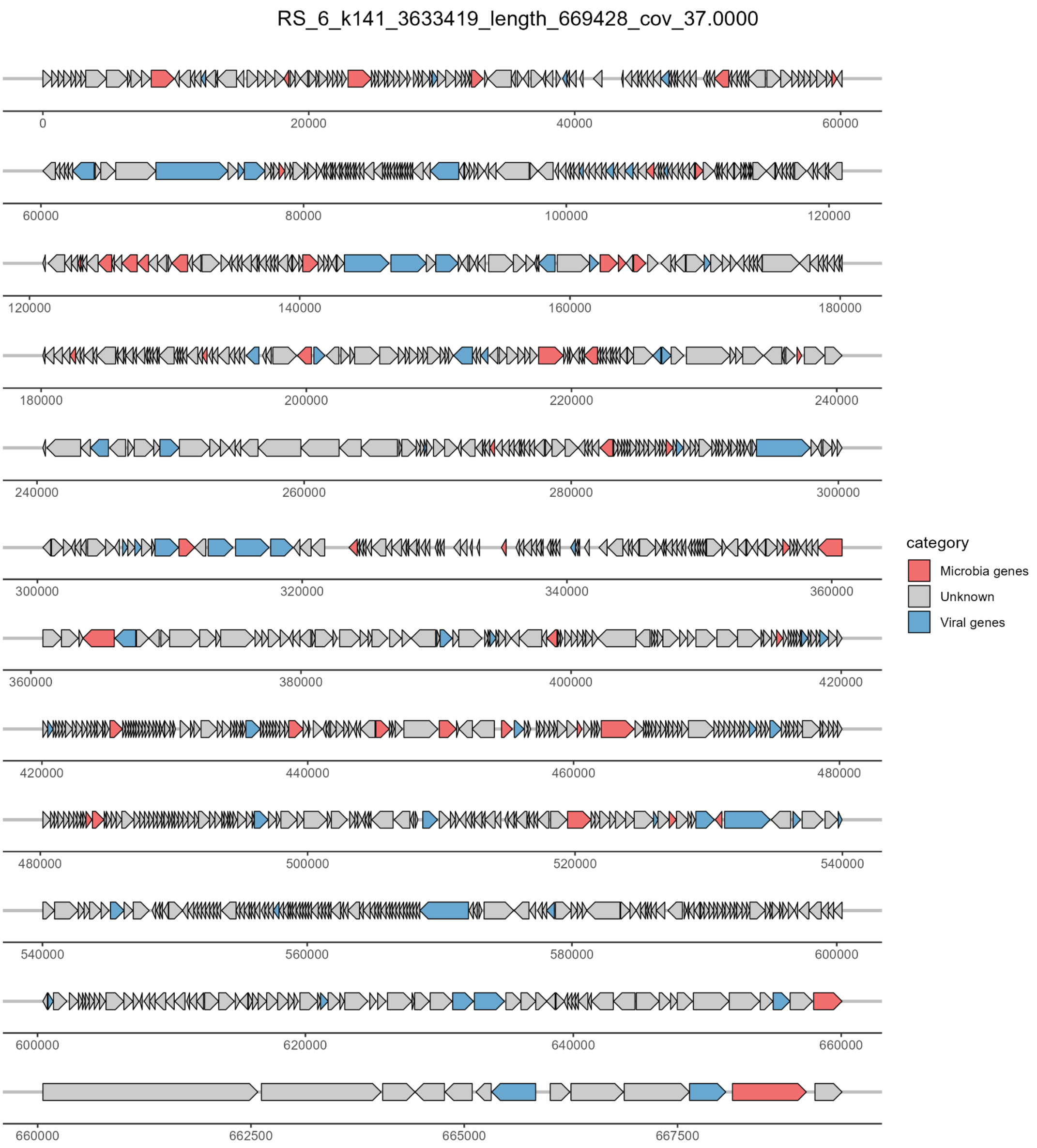

RS_6_k141_3633419_length_669428_cov_37.0000
category
Microbia genes
Unknown
Viral genes

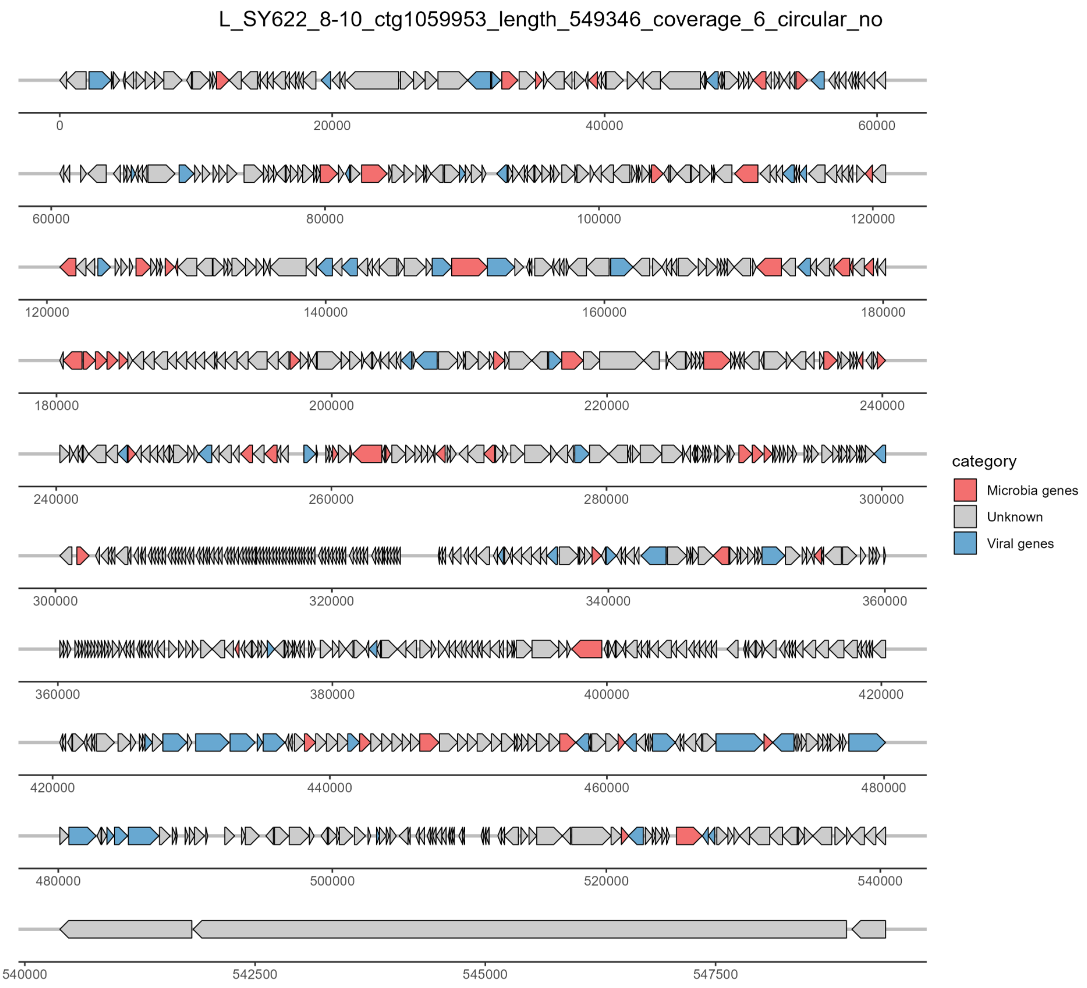


**Figure S25. Genome organization of the six largest viral genomes (>500 kb).** Predicted open reading frames are represented as arrows, oriented according to transcriptional direction. Genes are color-coded based on functional classification derived from CheckV annotation: genes with homology to microbial and viral genes are highlighted in red and blue, respectively.

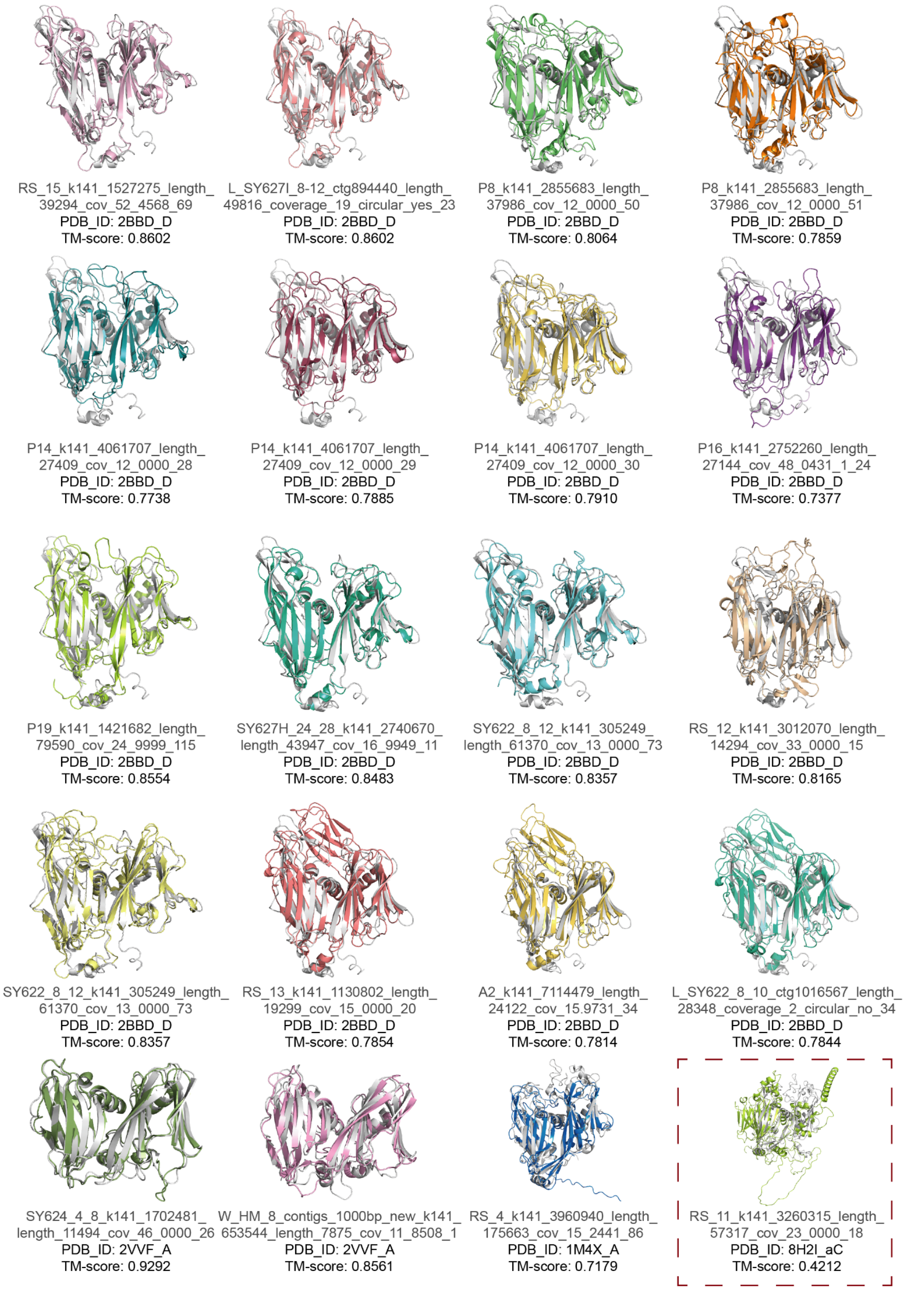


**Figure S26. Structural alignment of DJR-MCPs from the GCSV with reference proteins.** Predicted structures of 20 candidate DJR-MCPs from the GCSV were aligned with their corresponding reference structures. The DJR-MCPs predicted from the GCSV are displayed in color, while the corresponding reference structures from the PDB are shown in grey. Structural similarity was assessed using TM-align, and the PDB accession number and TM-score for each MCP-reference pair are indicated below the models. One MCP with a TM-score < 0.7 was considered non-homologous to the reference structure and is highlighted with a red dashed box.
